# Visualizing the impact of disease-associated mutations on G protein–nucleotide interactions

**DOI:** 10.1101/2024.01.30.578006

**Authors:** Kara Anazia, Lucien Koenekoop, Guillaume Ferré, Enzo Petracco, Hugo Gutiérrez-de-Teran, Matthew T. Eddy

**Affiliations:** Department of Chemistry; University of Florida; Gainesville, FL, 32611; USA; Department of Cell and Molecular Biology, Uppsala University; Uppsala, 75105; Sweden; Present address: Institut de Pharmacologie et de Biologie Structurale (IPBS), Université de Toulouse, CNRS, Université Toulouse III - Paul Sabatier (UT3), Toulouse, France; URD Agro-Biotechnologies Industrielles (ABI), CEBB, AgroParisTech, Pomacle, France

**Keywords:** GTPase, G protein, G protein-coupled receptor (GPCR), nuclear magnetic resonance (NMR), molecular dynamics (MD), cancer

## Abstract

Activation of G proteins stimulates ubiquitous intracellular signaling cascades essential for life processes. Under normal physiological conditions, nucleotide exchange is initiated upon the formation of complexes between a G protein and G protein-coupled receptor (GPCR), which facilitates exchange of bound GDP for GTP, subsequently dissociating the trimeric G protein into its Gα and Gβγ subunits. However, single point mutations in Gα circumvent nucleotide exchange regulated by GPCR–G protein interactions, leading to either loss-of-function or constitutive gain-of-function. Mutations in several Gα subtypes are closely linked to the development of multiple diseases, including several intractable cancers. We leveraged an integrative spectroscopic and computational approach to investigate the mechanisms by which seven of the most frequently observed clinically-relevant mutations in the α subunit of the stimulatory G protein result in functional changes. Variable temperature circular dichroism (CD) spectroscopy showed a bimodal distribution of thermal melting temperatures across all Gα_S_ variants. Modeling from molecular dynamics (MD) simulations established a correlation between observed thermal melting temperatures and structural changes caused by the mutations. Concurrently, saturation-transfer difference NMR (STD– NMR) highlighted variations in the interactions of Gα_S_ variants with bound nucleotides. MD simulations indicated that changes in local interactions within the nucleotide-binding pocket did not consistently align with global structural changes. This collective evidence suggests a multifaceted energy landscape, wherein each mutation may introduce distinct perturbations to the nucleotide-binding site and protein-protein interaction sites. Consequently, it underscores the importance of tailoring therapeutic strategies to address the unique challenges posed by individual mutations.

## Introduction

Signaling at the surface of eukaryotic cells, triggered by extracellular molecules such as neurotransmitters and hormones, begins with the interaction of G protein-coupled receptors (GPCRs) with intracellular G proteins. GPCR–G protein complex formation accelerates the exchange of bound GDP for GTP, dissociating the trimeric G protein complex into Gα and Gβγ subunits. Subsequently, each subunit activates intracellular signaling pathways until regulators of G protein signaling (RGS) proteins accelerate GTP hydrolysis, enabling the dissociated G protein subunits to reassemble and restart the process (1,2).

In typical physiological conditions, nucleotide exchange and the cyclical dissociation and reassociation of G proteins is carefully controlled through interactions of G proteins with partner receptor and RGS proteins. Single point mutations and residue-specific modifications within the Gα subunit of the trimeric G protein, whether acquired or inherited, subvert this process. Certain single point mutations within the Gα subunit accelerate GDP release or GTP uptake, causing constitutive activation and uncontrolled generation of secondary messengers (3,4) Other mutations prevent GDP release or dissociation of the Gα and Gβγ subunits, irreversibly halting secondary messenger production (5). In both cases, as G protein signaling is crucial to numerous physiological processes, circumventing regulation of G protein signaling is closely associated with the onset of a wide range of diseases (6–8), including inflammatory diseases (9), cardiovascular diseases (10,11), metabolic disorders (12,13), and cancers (14–17).

The occurrence of mutations in G proteins across a series of diseases has prompted intense investigation of the molecular mechanisms by which specific mutations in G proteins cause abnormal signaling. Determination of the structure of an R201C variant of the alpha subunit of the stimulatory G protein, Gα_S_, one of the most prevalent cancer-causing mutations in G proteins (18), revealed that this mutation caused constitutive activation of GDP-bound Gα_S_ by stabilizing an intramolecular hydrogen bond network (18). Integrative biophysical and biochemical experiments identified a triad of amino acids Gly203-Arg208-Glu245 (numbering refers to residues in Gα_I1_) that are conserved across G protein alpha subunits and connect GTP binding to the release of the Gβγ subunits from the trimeric complex (19); amino acid replacement of these residues prevented dissociation of the trimeric complex even with GTP bound to the alpha subunit (19). Integrative biochemical and structural analysis of a conserved catalytic glutamine residue revealed that different amino acid replacements of this conserved residue resulted in Gα variants with distinct structures and functions (20). Additional interesting biophysical studies demonstrated that Gα subunits contain pH-sensing residues, and mutation of these residues significantly changed the thermal stability and expression levels of the protein (21). Studies using NMR spectroscopy indicated that D150N, an oncogenic mutation in Gα_i_, favored an excited conformational state that did not tightly bind GDP (22). G protein structure-function relationships are complex, and despite the insights gleaned for several G protein variants, there remains a need for additional data on G protein variants, especially those associated with intractable diseases.

We leveraged an integrative experimental and computational biophysical approach to investigate seven different mutations in Gα_S_ and their influence on Gα_S_ structure-function relationships. While our study focused on Gα_S_, as it is one of the more frequently mutated G protein α subunits observed across multiple cancers (17,23), many of the mutations investigated here are also present in other G proteins. Variable-temperature circular dichroism (CD) spectroscopy was utilized to compare the thermal melting behavior and related global structural features of Gα_S_ and Gα_S_ variants. In complementary molecular dynamics (MD) simulations, we observed changes in hydrogen bonding patterns among the variants consistent with experimental thermal melting data, providing structural interpretation of observed differences in melting temperatures. Additionally, we employed saturation-transfer difference NMR (STD-NMR) spectroscopy in aqueous solutions to examine how mutations impacted the interaction of nucleotides with residues in the nucleotide binding pocket and provide a structural and dynamic interpretation of this data. The integration of these complementary data sets provided a view of the impact of mutations both at the global structural level and local nucleotide binding pocket of Gα_S_ variants, which we discuss in the context of available kinetics and cell signaling data.

## Results

### Preparation of functional human G*α*_S_ and G*α*_S_ variants

We compared structure-function relationships of Gα_S_ and seven Gα_S_ variants containing single amino acid replacements: R201C, G226A, Q227L, R228C, R258A, R265H, and A366S (Figure 1; Table S1). The positions of the selected mutations are located across multiple distinct regions within the Gα_S_ structure that undergo conformational changes upon nucleotide exchange, namely the Switch I, Switch II, and Switch III regions (24). Selection of Gα_S_ residue locations for this study was designed to survey all three structural regions. R201 is located in Switch I; G226, Q227, and R228 are located in Switch II; R258 and R265 are located in Switch III (Figure 1). Additionally, A366 is located between the β5 and α5 regions adjacent to the nucleotide binding pocket, approximately 3.5 to 4.0 Å from the imidazole group of the guanine ring system. To provide an internal control, the study also included “mini-Gα_S_”, an engineered G protein with a truncation of the alpha helical domain and multiple mutations introduced to facilitate its crystallization in complex with GPCRs (25,26). Mini-Gα_S_ recapitulates certain essential aspects of Gα_S_ binding to GPCRs, however it does not bind nucleotides with the same affinity as Gα_S_ and does not undergo global conformational rearrangements upon nucleotide exchange (25).

**Figure 1.**
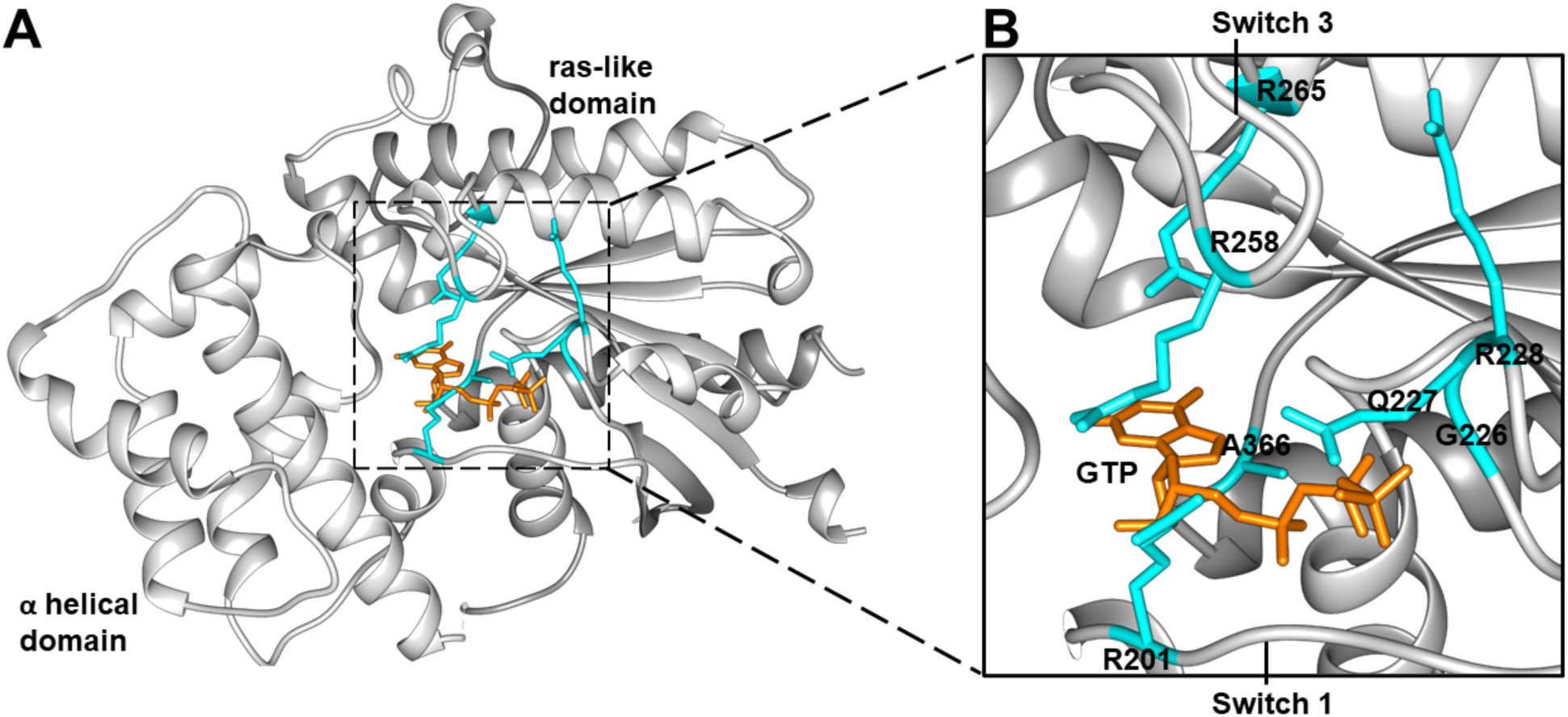
Crystal structure of Gα_S_ with annotated disease-associated mutations. *A,* the GTP-bound Gα_S_ crystal structure (PDB: 1AZT)^1^. GTP is shown in orange stick representation and residues selected for investigation shown in blue stick representation. *B*, expanded view from the dashed box shown in *A*. GTP, disease-associated residues, and the Switch 1 and Switch 3 structural regions are annotated.

The impacts of the seven mutations on G protein signaling have been documented in earlier studies (Table S1). Three different gain-of-function mutations cause increased cyclic AMP (cAMP) accumulation even in the absence of GPCR stimulation, which include Gα_S_[R201C] (18), Gα_S_[Q227L] (27,28), and Gα_S_[A366S] (4). Four different loss-of-function mutations result in decreased cAMP production, which include Gα_S_[G226A] (5), Gα_S_[R228C] (18), Gα_S_[R258A] (18) and Gα_S_[R265H] (18).

Human Gα_S_ and all Gα_S_ variants were expressed in *E.coli* and purified to >95% homogeneity using protocols adapted from earlier studies (18,25); see Experimental Procedures and Figure S1*A–B* for additional details. CD spectra of purified Gα_S_ and all Gα_S_ variants showed minima at 210 and 220 nm (Figure 2*A*, S2), typical of folded proteins containing alpha helical secondary structure, confirming all samples were folded. For Gα_S_, structural rearrangements in the three switch regions associated with nucleotide exchange result in an increase in intrinsic tryptophan fluorescence (24). To verify protein function, we monitored changes in intrinsic tryptophan fluorescence upon exchange of GDP for the non-hydrolysable GTP analog GTPγS (Figure S1). Rates of tryptophan fluorescence enhancement for Gα_S_ and Gα_S_ variants were consistent with values reported in prior studies (18). Some mutations, such as G226A, resulted in decreased fluorescence, rather than an increase, but this too was consistent with literature data (5).

**Figure 2.**
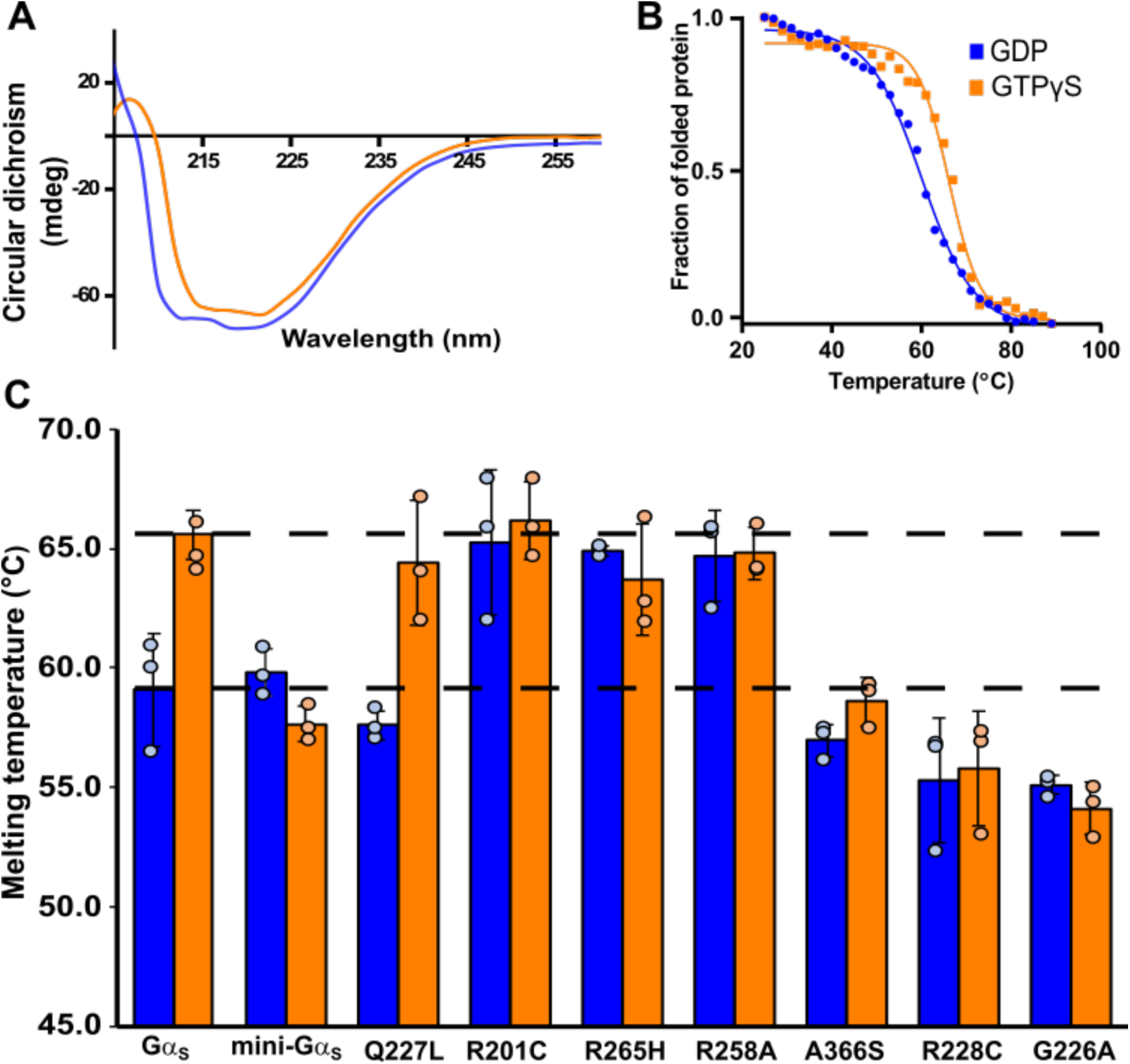
Thermal melting temperatures of Gα_S_ and Gα_S_ variants determined by variable-temperature circular dichroism spectroscopy. *A,* representative CD spectrum of Gα_S_ bound to GDP (blue curve) and bound to GTPγS (orange curve). *B,* thermal unfolding of Gα_S_ bound to GDP or GTPγS monitored by variable temperature single wavelength CD spectroscopy. Same color scheme as *A*. *C,* histogram of melting temperatures determined from fitting the variable temperature CD data of Gα_S,_ mini-Gα_S_ and Gα_S_ variants obtained in complexes with GDP or GTPγS. The dashed horizontal lines are located at the mean T_m_ values for Gα_S_ in complexes with GDP or GTPγS. Error bars were determined from the standard deviation of triplicate measurements.

### Disease-associated mutations alter the thermal stability of G*α*_S_ complexes with nucleotides

Previous investigations documented that Gα_i1_ complexes with GTP or GTP analogs exhibited a ~10 °C higher thermal denaturation temperature compared with GDP-bound Gα_i1_ (19,21,29). These observations were interpreted to indicate that Gα_i1_ adopted different conformations between the GDP and GTP-bound states and that each state exhibits a different global structural stability (19,21,29).

We compared the thermal unfolding of Gα_S_, the seven Gα_S_ variants, and mini-Gα_S_ using variable temperature CD spectroscopy in the presence of either GDP or GTPγS. The unfolding temperature (T_m_) of GTPγS-bound Gα_S_ was observed to be ~9 °C higher than GDP-bound Gα_S_ (Figure 2*B*, 2*C*), consistent with earlier findings of Gα_i1_ (19,21,29). In contrast, for all Gα_S_ variants, with the exception of Gα_S_[Q227L], we observed at most only minor differences in the T_m_ values between GDP-bound and GTPγS-bound samples (Figure 2*C*).

The melting temperatures of the Gα_S_ variants Gα_S_[R228C], Gα_S_[G226A], and Gα_S_[A366S] bound to either GDP or GTPγS were found to be comparable to the observed melting temperature of GDP-bound Gα_S_ (Figure 2C). Based on these findings, we infer that these three variants do not exhibit distinct global conformations in the presence of GDP or GTP. Instead, they adopt a global conformation more closely aligned with GDP-bound Gα_S_, even when exposed to a saturating concentration of GTPγS. In contrast, the variants Gα_S_[R201C], Gα_S_[R265H] and Gα_S_[R258A] showed relatively higher melting temperatures in the presence of both GTPγS and GDP, falling within a range similar to Gα_S_ bound to GTP (T_m_=65.2±1.5°C) (Figure 2*C*). These observations indicate that these three variants do not exhibit distinct global conformations in the presence of GDP or GTP and instead adopt a global conformation more closely aligned with GTP-bound Gα_S_ even in the presence of GDP. A noteworthy exception to this altered thermal response was observed in Gα_S_[Q227L], which exhibited significant differences in melting temperatures between GDP-bound and GTPγS-bound samples, resembling the thermal behavior of Gα_S_ (Figure 2C, S2).

For comparison, we also recorded thermal unfolding data for mini-Gα_S_. As expected, we observed at most minimal differences in temperature between mini-Gα_S_ in the presence of GDP or GTPγS (Figure 2), in line with the absence of structural differences observed in the presence of GDP or GTPγS and consistent with reports that claimed mini-Gα_S_ did not respond to the presence of GTP(25).

To further investigate the interplay between mutations and bound nucleotides on the thermal melting profiles of Gα_S_ and Gα_S_ variants, we recorded variable temperature CD spectra of Gα_S_ and selected variants that exhibited higher T_m_ values prepared without nucleotide added during the protein purification process (Figure S3). Gα_S_, Gα_S_[R201C], Gα_S_[R258A], and Gα_S_[R265H] could be expressed and isolated without nucleotide added during the purification process. Apo Gα_S_ exhibited a T_m_ of ~62 °C, higher than the T_m_ of GDP-bound Gα_S_ and lower than the T_m_ of GTP-bound Gα_S_ (Figure S3). This suggests apo Gα_S_ adopts a global conformation that differs from both GDP-bound and GTP-bound Gα_S_. Interestingly, although Gα_S_[R201C], Gα_S_[R258A], and Gα_S_[R265H] all exhibited higher T_m_ values for both complexes with GDP, we observed strikingly different behavior for apo preparations of these samples. For apo Gα_S_[R201C], we observed a lower T_m_ value of around 59 °C. Notably, both apo Gα_S_[R258A] and Gα_S_[R265H] exhibited higher T_m_ values similar to the T_m_ values in the presence of either nucleotide for these variants (Figure S3). This suggests that the global conformations of Gα_S_[R258A] and Gα_S_[R265H] are similar both in the presence and absence of nucleotides. However, Gα_S_[R201C] appears to adopt different conformations in the absence or presence of nucleotides.

### MD simulations reveal changes in hydrogen bonding that correlate with thermal melting profiles of different *Gα_S_* variants

To investigate a structural basis for the observed impacts of mutations on the melting temperatures of Gα_S_ variants, we conducted molecular dynamics (MD) simulations on Gα_S_ and several Gα_S_ variants in complexes with GDP and GTP. Because detailed, atomistic simulations are time and resource intensive, we focused our efforts on simulating Gα_S_ and the variants Gα_S_[R228C], Gα_S_[Q227L] and Gα_S_[R258A]. These variants were selected because they each exhibited distinct thermal melting profiles in variable-temperature CD experiments (Figure 2). As discussed above, Gα_S_[R228C] exhibited a lower T_m_ for both GDP and GTP-bound samples, similar to the thermal melting profiles of Gα_S_[A366S] and Gα_S_[G226A]. Gα_S_[R258A] exhibited a higher T_m_ for both GDP and GTP-bound samples, paralleling the thermal melting profiles of Gα_S_[R201C] and Gα_S_[R265H]. Gα_S_[Q227L] was the only variant that showed an increase in T_m_ between GTP and GDP-bound samples, as in the case of Gα_S_. In earlier studies of Gα_i1_, a higher T_m_ was observed for GTP-bound Gα_I1_, which was attributed to increased non-covalent interactions between Switch I and Switch II and between Switch I and Switch III upon complex formation of Gα_i1_ with GTP or GTP analogs (19,20). From this earlier work, we hypothesized that the thermal melting behavior of Gα_S_[R228C], Gα_S_[Q227L] and Gα_S_[R258A] should thus correlate with different patterns of non-covalent interactions within the Switch regions between these variants.

Following this line of reasoning, we analyzed the occurrence of hydrogen-bonding between the switch regions within both GTP-bound and GDP-bound MD simulations of Gα_S_ and the three variants. Analysis of hydrogen bonding interactions captured from MD simulations revealed that Gα_S_[R228C] is the only variant where an interaction between Ser205 in Switch I and Glu230 in Switch II is absent for the GTP-bound conformation (Figure 3). This suggests that this interaction may at least be partially related to maintaining structural integrity of the GTP-bound protein and absence of this interaction in Gα_S_[R228C] may partially explain its observed lower thermal stability. Further, we also observed two additional interactions between Switch II and Switch III that were present for GTP-bound Gα_S_ that were absent for Gα_S_[R228C] bound to GTP between Arg228 and Glu259 and between Arg228 and Glu268 (Figure 3). Comparing all GTP bound proteins, we observed the fewest number of interactions between Switch regions for Gα_S_[R228C], consistent with its observed lower thermal stability.

**Figure 3.**
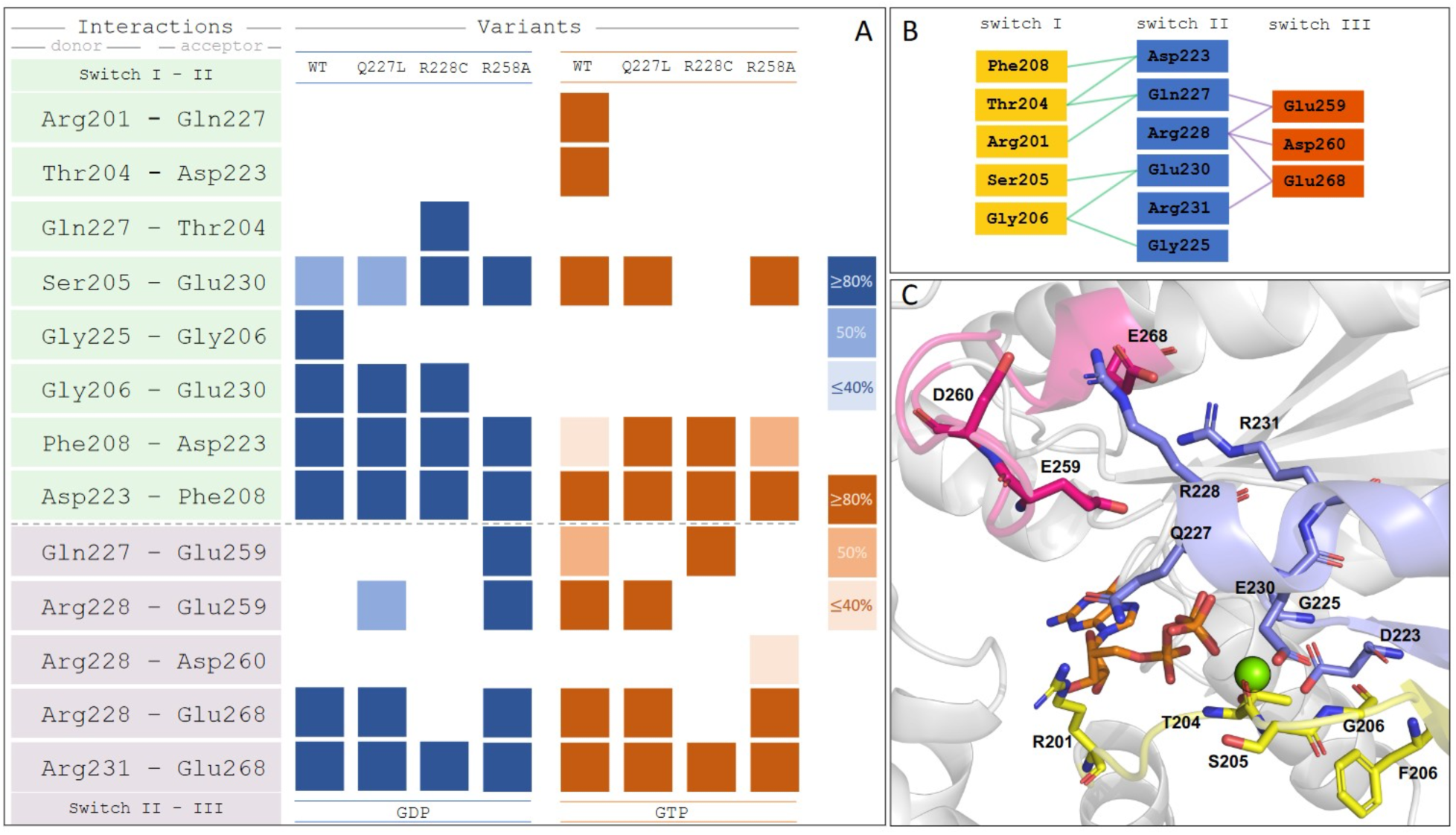
Analysis of hydrogen bond interactions between switch regions extracted from MD simulations. *A*, hydrogen-bond interaction profiles for Gα_S_ (labeled ‘WT’), Gα_S_[Q227L], Gα_S_[R228C], and Gα_S_[R258A], focused on interactions between Switch I, Switch II, and Switch III regions. Colored boxes indicate the existence of a persistent hydrogen-bond, determined with different thresholds of occurrence along the given MD simulation, proportional to the color intensity for GDP-bound (blue) or GTP-bound (orange) proteins. *B*, schematic representation of all hydrogen-bond interactions depicted in the H-bond matrices. *C*, the structure of GTP-bound Gα_S_ with the involved residues from the analysis annotated. Residues from Switch I are colored yellow, Switch II colored blue and Switch III colored red.

Analysis of Gα_S_[R258A] bound to GDP, which showed a marked increase in thermal stability compared with Gα_S_, revealed increased hydrogen bond interactions between Switch II and Switch III and one fewer interaction between Switch I and Switch II (Figure 3), including the presence of two interactions absent for Gα_S_, between Gln227 and Glu259 and between Arg228 and Glu259 (Figure 3). Interestingly, Gα_S_[R258A] is the only GDP-bound variant where one interaction between Switch I and Switch II is absent, Gly206-Glu230, which is also absent for GTP-bound Gα_S_ and all GTP-bound variants, a possible additional indicator of increased thermal stability for Gα_S_[R258A].

From the MD simulations, we also compared the dynamics of each switch region, described by the root mean square fluctuation (RMSF), among Gα_S_ and the different Gα_S_ variants considered in this analysis. RMSF values were calculated from averaging over all MD trajectories for each variant and nucleotide combination. Fluctuations were determined as the average backbone deviation relative to the average position of the full trajectory per residue. We observed the largest RMSF for Gα_S_[R228C] for both the GDP and GTP–bound forms of this variant (Figure S4), particularly around residues 225 to 230 of Switch II. We also observed increased RMSF values across five residues of Switch III for the same variant (Figure S4). This appears consistent with decreased hydrogen bonding observed for residues between Switch II and Switch III of Gα_S_[R228A] (Figure 3).

### STD-NMR spectroscopy with Gα_S_ and Gα_S_ variants in complexes with nucleotides

While CD data provided information on global structure and stability of Gα_S_ and Gα_S_ variants, we also aimed to investigate how mutations potentially altered local interactions between the nucleotides and Gα_S_ within the nucleotide binding pocket. Saturation transfer difference (STD)-NMR spectroscopy has been employed extensively to provide information on protein-ligand interactions and is especially powerful for elucidating information on moderately-to-weakly binding ligands (K_d_ > 1µM) (30–34). Well-isolated resonances from the protein, such as methyl groups that exhibit unique chemical shifts, are saturated in the ^1^H-NMR spectrum. If a ligand interacts with the protein, polarization is transferred to the ligand from the protein via spin diffusion (31). By subtracting the saturated spectrum from a reference spectrum, a difference spectrum is obtained where signals from protons of the interacting ligand are predominantly observed. The intensities of the observed ^1^H ligand signals depend on both the average distance and the dynamics of non-covalent interactions between the protein and ligand (31,33).

We recorded one-dimensional ^1^H STD-NMR data with Gα_S_ and the seven Gα_S_ variants in the presence of a 50-fold excess of GDP or GppNHp (see Experimental procedures for more details). GppNHp was used rather than GTPγS due to different availabilities of the two compounds at the time of the study. Both GppNHp and GTPγS are widely used in studies of G proteins as both have been shown to exhibit affinities for most G proteins that differ by less than an order of magnitude (35,36). ^1^H NMR signals for both GDP and GppNHp were identified and assigned in one-dimensional reference spectra (Figure S5) by transferring assignments reported from earlier studies (Biological Magnetic Resonance Bank accession code bmse000270) (37). From exploratory STD-NMR experiments, we identified four ^1^H signals that were well-resolved and free from residual protein background signals that were assigned to protons in the guanosine and sugar ribose groups, labeled as protons ‘a’ through ‘d’ (Figure S5). STD-NMR experiments with GDP utilized signals from these four protons; in experiments with GppNHp, protons ‘c’ and ‘d’ were overlapped with background signal from the protein and we therefore excluded them from analysis of samples containing GppNHp. Examination of the structure of Gα_S_ (PDB ID 1AZT) showed multiple residues within ~6 Å of the protons monitored in STD-NMR experiments (Figure S6). We anticipated these residues would make the largest contributions to observed STD-NMR signals and predicted that either structural rearrangement of these residues or their fluctuation relative to the bound nucleotide would result in changes in observed STD-NMR signals.

In initial STD-NMR experiments with Gα_S_ and GDP, we optimized the experimental signal-to-noise by varying the saturation transfer time from 0.5 to 3.0 seconds (Figure S7). These data confirmed that interactions between nucleotides and Gα_S_ were detected in STD-NMR experiments and allowed optimization of the transfer time. We observed a significant increase in the STD-NMR signal up to 2.0 seconds, and only minor increases in STD-NMR signals were observed with a 3.0 second transfer time. We therefore selected 2.0 seconds for all subsequent experiments. Control experiments recorded for samples containing either only Gα_S_ and buffer or only nucleotide and buffer showed no STD-NMR signals, as expected (Figure 4). This confirmed that signals observed in STD-NMR experiments were due to specific interactions between nucleotides and Gα_S_ or Gα_S_ variants.

**Figure 4.**
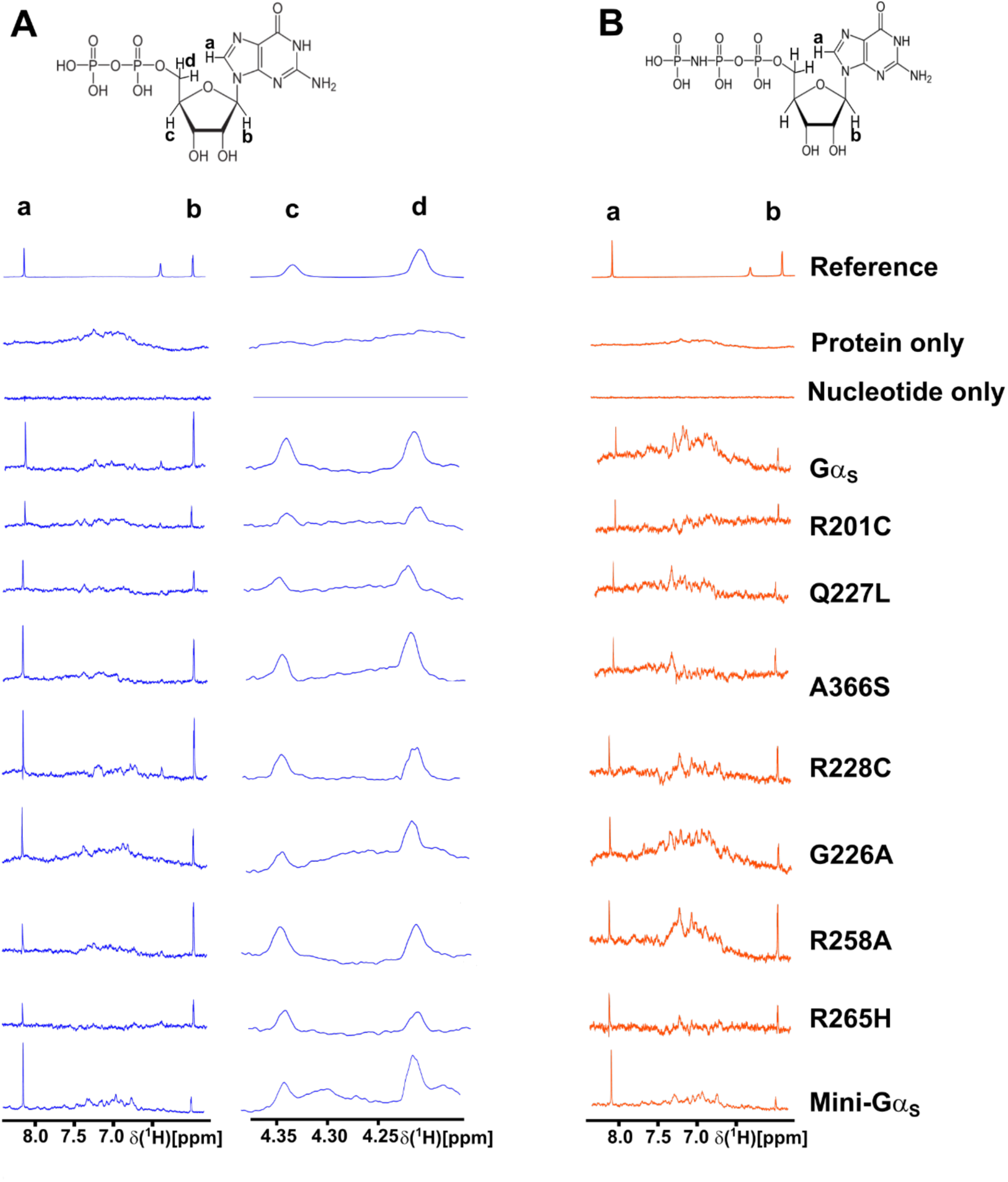
One-dimensional ^1^H STD-NMR spectra of Gα_S_ and Gα_S_ variants in complexes with GDP and GppNHp. *A,* the chemical structure of GDP and 1D STD-NMR spectra of complexes with GDP. Protons observed in the STD-NMR spectra are annotated ‘a’ to ‘d’. Below the chemical structure are expanded regions from 1D STD-NMR spectra shown in Figure S8 containing the annotated signals. “Reference” is a 1D ^1^H NMR spectrum of GDP, “protein only” is an STD-NMR control experiment with a sample containing Gα_S_ and buffer but no nucleotide, and “nucleotide only” is a STD-NMR control experiment with a sample containing nucleotide and buffer but no protein. *B,* the chemical structure of GppNHp and 1D STD-NMR spectra of complexes with GppNHp. Other presentation details are the same as in *A.* Views of the presented NMR data were expanded from the full spectra shown in Figure S7.

In STD-NMR spectra of Gα_S_ or Gα_S_ variants with GDP, we observed significant variation in the signal intensities of protons ‘a’ through ‘d’ (Figure 4 and Figure S8). Mini-Gα_S_ in particular showed the smallest signal intensities in STD-NMR spectra with respect to Gα_S_ for protons ‘b’ through ‘d’, which appears consistent with its expected weaker nucleotide binding. In STD-NMR spectra of Gα_S_ or Gα_S_ variants with GppNHp, we also observed a large variation in signal intensities, with mini-Gα_S_ again showing the weakest signal intensities (Figure 4 and Figure S8). We integrated the observed signal intensities from the STD-NMR spectra and used them to calculate corresponding STD amplification factors (STD-AF) by multiplying the observed STD-NMR signal by the molar excess of ligand over protein (see Experimental Procedures), following previously validated protocols (30,31). The STD-AF converts the STD signal intensity, which depends on the fraction of bound ligand, to a value that depends on the fraction of bound protein (30,31). We then calculated and compared the STD-AF values for all samples of Gα_S_ and Gα_S_ variants with GDP and GppNHp (Figure 5). The STD-AF values were normalized with respect to the ‘a’ proton of the guanine nucleobase, because in MD simulations, the immediate environment around the ‘a’ proton showed very little change (see below and Figure S9).

**Figure 5.**
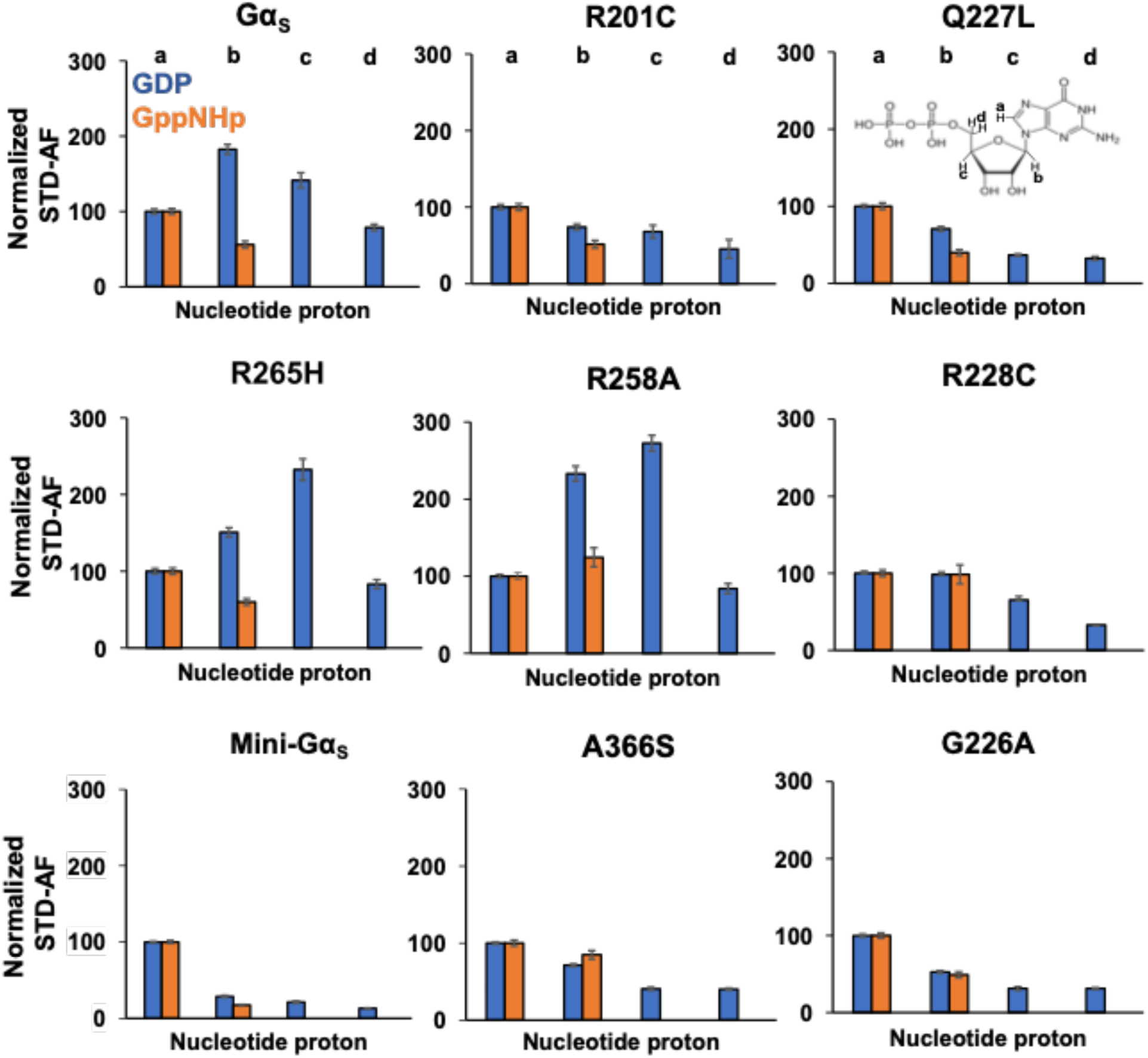
Normalized STD amplification factors measured for GDP and GppNHp in complex with Gα_S_ or Gα_S_ variants. Normalized STD amplification factors (STD-AF) determined for GDP (blue) and GppNHp (orange) in complex with Gα_S_ or Gα_S_ variants. The specific protons used to quantify the STD-AF values are annotated on the chemical structure of GDP. STD amplification factors were normalized to the ‘a’ proton. Error bars were determined from the signal-to-noise ratios in the corresponding NMR spectrum.

The normalized STD-AF values for samples of Gα_S_ or Gα_S_ variants prepared with either GDP or GTP provided a “fingerprint” of the extent of interaction of the nucleotide protons with local protons from the protein. For Gα_S_, relative to the ‘a’ proton, we observed an increased STD-AF for the b and c protons with GDP, and a decrease in the STD-AF for the ‘b’ proton with GppNHp (Figure 5 and Figure 6). For five of the Gα_S_ variants, we observed the opposite effect for samples prepared with GDP, specifically we observed decreasing STD-AF for protons ‘b’, ‘c’, and ‘d’. We observed the smallest STD-AF values for both GDP and GppNHp for mini-Gα_S_. Though the STD-AF values do not necessarily directly reflect binding affinities, the lower STD-AF values observed for mini-Gα_S_ are consistent with our expectations that this engineered protein exhibits significantly altered binding with nucleotides (25). Two variants exhibited an opposite trend: both Gα_S_[R258A] and Gα_S_[R265H] showed more intense STD-AF values for ‘b’ and ‘c’ protons for complexes with GDP, indicating increased interactions with GDP within the nucleotide binding pocket (Figure 6). For protein complexes with GppNHp, STD-AF values calculated for ‘a’ and ‘b’ protons show a decrease in the value of the ‘b’ proton relative to ‘a’ for Gα_S_ (Figure 5 and Figure 6), establishing a ‘fingerprint’ of interaction for the native protein. Compared with the STD-AF profile of Gα_S_, the STD-AF values for five of the Gα_S_ showed similar profiles, with two exceptions. Both Gα_S_[R258A] and Gα_S_[R228C] showed larger values observed for the ‘b’ proton for their complexes with GppNHp (Figure 5 and Figure 6). While we did observe an STD-NMR signal for GppNHp in the presence of mini-Gα_S_, the value for the ‘b’ proton was the lowest observed within the study, consistent with the expectation of altered binding of this engineered protein to the GTP analog (Figure 5 and Figure 6).

**Figure 6.**
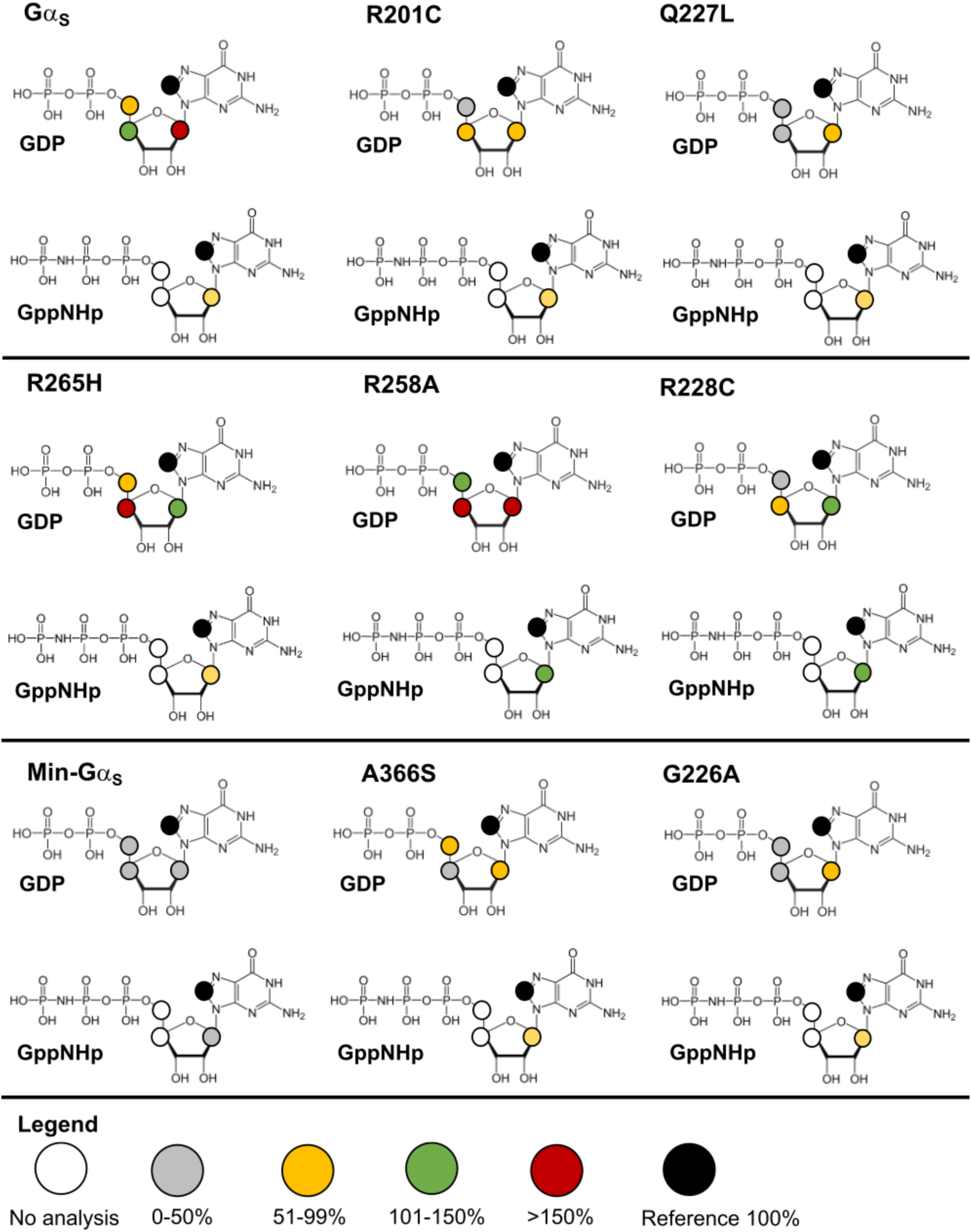
STD-NMR amplification factors mapped onto the chemical structures of GDP and GppNHp for complexes with Gα_S_ and Gα_S_ variants. For each proton observed in the STD-NMR spectra, the relative STD-AFs are colored as a percentage of the STD-AF normalized with respect to the ‘a’ proton (same as in Figure 3 and 4) indicated with the black circle. The STD-AFs are colored according to their relative intensities compared to those for the ‘a’ proton: relatively weaker interactions representing < 50% (grey), weak-moderate interactions representing 51% to 99% (yellow), moderate interactions representing 101% to 150% (green) and stronger interactions representing > 150% (red). The white circles indicate that no analysis was performed for those protons due to overlap of nucleotide and background residual protein signals.

### Structural characterization of changes around the nucleotide binding pocket from MD simulations

Observations from the STD-NMR data reflect changes in the local structure and dynamics of amino acids within the nucleotide binding pocket of Gα_S_ and Gα_S_ variants; however, different configurations of the nucleotide binding region may result in similar STD-AF values, providing a more granular view. To obtain a more detailed view, we analyzed our MD simulations to investigate potential differences in the dynamics and configuration of the binding pocket, focusing on GDP-bound and GTP-bound Gα_S_ and the three variants explored in the earlier structural investigations underlying different thermal melting behavior, Gα_S_[R228C], Gα_S_[Q227L] and Gα_S_[R258A]. For each protein and for each nucleotide complex, we monitored each of the four protons on the nucleotide, ‘a’ through ‘d’, observed in the STD-NMR experiments over the course of the simulations. In each simulation, we identified the closest protons in the protein within a radius of ~6Å of the nucleotide and the calculated the duration that each residue spends in proximity to one or more of the monitored protons (Figure 7 and Figure S9).

**Figure 7.**
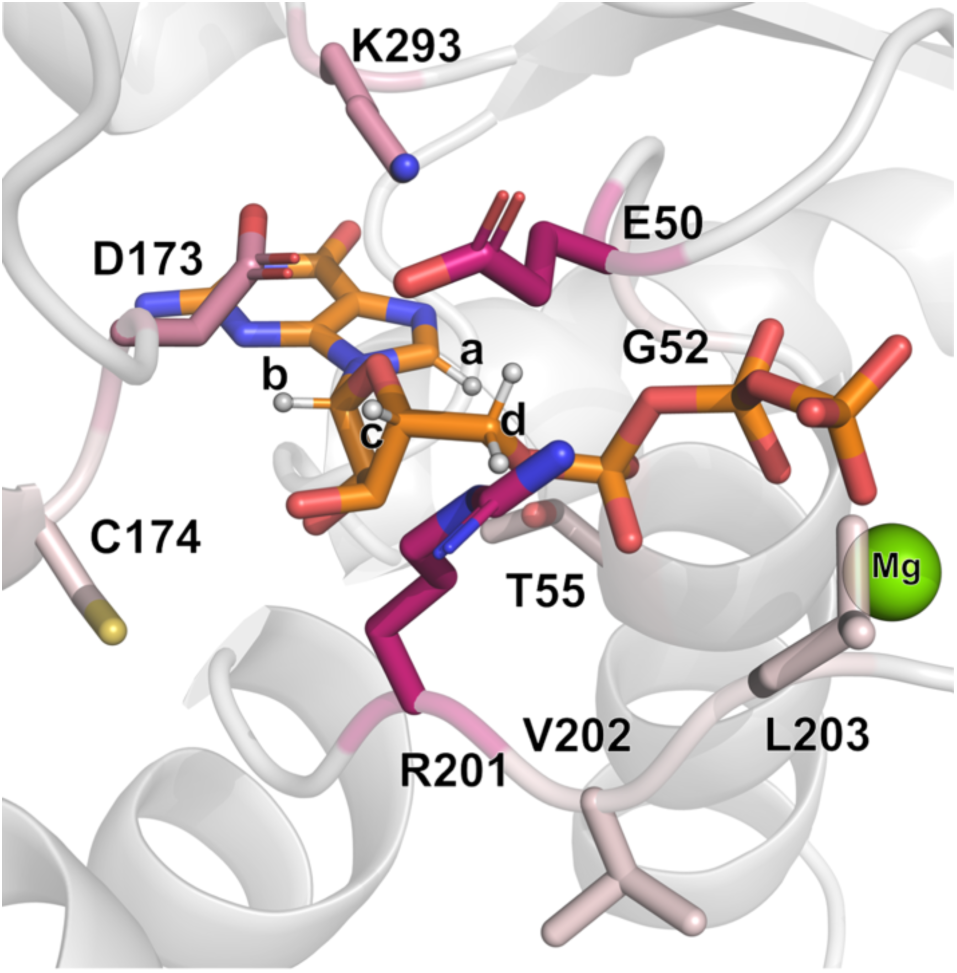
Variation in the structure and dynamics of the nucleotide binding pocket among Gα_S_ and Gα_S_ variants. Nucleotide binding pocket of Gα_S_ bound to GTP (shown in orange stick representation, PDB:1AZT) with protons seen in STD-NMR interactions annotated ‘a’ through ‘d’. The color palette for the residues interacting with these protons denotes increasing shades of pink which are proportional to the frequency of the residue as being the closest to the given proton, with darker shades of pink indicating higher frequencies as a function of the mutations explored.

We observed both similarities and differences in the local environment among Gα_S_ and the three Gα_S_ variants. The ‘a’ proton on the guanine ring of the nucleotide is in persistent proximity with G52 for both GDP-bound and GTP-bound Gα_S_ and for all variants, with an overall mean occupancy across all proteins of 97.4±1.4% (Figure 7 and Figure S9). The associated standard error of the mean (s.e.m.) was used as an indicator of the variability, which was relatively low in this case. As discussed above, this mostly uniform local environment around proton ‘a’ rationalized our normalization of the STD-AF values with respect to its signal intensity (Figure 5 and Figure 6).

The ‘b’ proton is mainly interacting with Asp173 (mean occupancy of 67±14%) and Lys293 (32.2±14.1%, Figure S9). Most of this variability is due to the significant difference between the GDP and GTP-bound forms in both Gα_S_[Q227L] and Gα_S_[R258A] mutants, where mainly the GTP-bound form shows an even contact between the ‘b’ proton and Asp173 and Lys293. Interestingly, the STD-AF normalized values of these two mutants show a gap for the ‘b’ proton interactions between their corresponding GDP and GTP. For the remaining protons, we do not have measurement of the GTP-bound normalized STD-AF values. Nevertheless, we can examine results from MD simulations to make predictions on their behavior. Proton ‘c’ exhibits the highest variability in the identity of the closest residue, among the different Gα_S_ forms, with a more or less shared interaction with Arg201, Asp173 and Glu50, exception made for the Gα_S_[Q227L] form (where it almost exclusively contacts with Glu50 when bound to GDP or with Arg201 when bound to GTP). Analysis of the ‘d’ protons in GDP was more complex since this signal is due to two equivalent protons. To address this, we analyzed the arithmetic average of the independent measurements for each proton, and the results show a very constant and even distribution of the closest interacting residues, with Glu50 and Arg201 making an even interaction with the ‘d’ protons in all simulations. Moreover, in all four protein systems (Gα_S_ or variants Gα_S_[Q227L], Gα_S_[R258A] and Gα_S_[R228C]) the GTP-bound form displays a distinct interaction of the ‘d’ protons with Gly52 (average occupancy 13.0±3.8%, vs 0.6±0.8% in the corresponding GDP-bound forms, Figure S9), suggesting a different microenvironment of these protons as a function of the nucleotide bound.

## Discussion

The current study presents complementary approaches to dissect the complex structure-function relationship of Gα_S_ and the impact of single-point mutations on Gα_S_ structure, dynamics, and function. Comparison of variable temperature CD data of Gα_S_ with Gα_S_ variants in complex with GDP or a GTP analog show striking differences (Figure 2). Three mutations, A366S, R228C, and G226A showed reduced thermal stability for the GTP-bound protein, comparable to the T_m_ for the same proteins in complex with GDP (Figure 2). These results indicated that these mutations impaired the ability of the protein to adopt a conformation observed for Gα_S_ in complex with GTP. This appears in line with earlier studies of Gα_I1_ that showed replacement of key residues in a conserved ‘Gly-Arg-Glu’ triad reduced protein stability, which was attributed to loss of the ability of Gα_i1_ to undergo conformational changes upon interacting with GTP (19). The mini-Gα_S_ protein also exhibited decreased T_m_ in the presence of GTPγS, which was expected because this engineered protein lacks several segments of the switch loops that confer stability and function to Gα_S_. While all three mutants exhibit reduced thermal stability for GTP-bound protein, G226A and R228C are associated with decreased cAMP production (18) while A366S is associated with increased cAMP production (4). Both G226A and R228C are in Switch II whereas A366S is located near the nucleotide binding pocket, where this residue coordinates interactions between the nucleotide and β6–α5 loop (Figure 1). The mutation corresponding to Gα_S_[G226A] in Gα_i_ and was observed to disrupt Mg^2+^ binding and inhibit structural changes in the presence of GTP (5). The Ser366 in Gα_S_[A366S] has been shown to weaken affinity for GTPγS by sterically crowding the nucleotide binding pocket (38). Thus, while Gα_S_[G226A] and Gα_S_[A366S] share similar thermal melting responses to different nucleotides, the different mechanisms by which each mutation impacts function results in distinct functional changes.

Gα_S_[R201C], Gα_S_[R258A], and Gα_S_[R265H] all exhibit increased thermal stability for complexes with GDP, similar to the T_m_ measured for Gα_S_ in complex with GTPγS (Figure 2). This indicates that additional hydrogen bonds present in GTP-bound Gα_S_ are not broken for these variants in complex with GDP. This observation is consistent with the structure of Gα_S_[R201C] in complex with GDP (PDB:6AU6), which shares key features of the structure of Gα_S_ bound to GTP (18). A distinguishing feature among Gα_S_[R201C], Gα_S_[R258A], and Gα_S_[R265H] is the thermal melting profile of the apo proteins (Figure S3). The lower thermal melting temperature for apo Gα_S_[R201C], suggests that binding either GDP or GTP is required to form stabilizing interactions between the switch regions. In contrast, the higher thermal melting temperatures for apo Gα_S_[R258A], and apo Gα_S_[R265H] suggest that those stabilizing interactions are present even in the absence of any bound nucleotide. Based on this comparison, we predict the structures of apo Gα_S_[R258A] and Gα_S_[R265H] will be highly similar and will also differ from the structure of apo Gα_S_[R201C]. Differences in the behavior of the apo proteins appear to correlate with the impact of each mutation on function. While Gα_S_[R258A] and Gα_S_[R265H] exhibit an increased GTP binding rate and decreased GDP dissociation rate (18,39), R201C exhibits the opposite behavior (18).

Surprisingly, Gα_S_[Q227L] is the only variant that exhibited a difference of T_m_ values between complexes with GDP and GTPγS, similar to the thermal melting behavior of Gα_S_ (Figure 2). This suggests that this particular mutation does not inhibit the protein’s ability to adopt different global conformations for complexes with different nucleotides. This is unexpected because this mutation also increases constitutive cAMP production and exhibits reduced GDP dissociation and reduced GTP uptake (40). In MD simulations, we observed that two interactions between Switch I and Switch II present in Gα_S_, Arg201–Gln227 and Thr204–Asn223, were absent in not only R228C and R258A but also Q227L, suggesting that these interactions are not essential for the stability of the Gα_S_ complex with GTP (Figure 3). Gα_S_[Q227L] was the only variant where an interaction between Arg228 and Glu259 between Switch II and Switch III is preserved for the GTP-bound protein, as also observed for the native Gα_S_ (Figure 3). This feature may at least partially rationalize the observed correlation in similar thermal melting behaviors of Gα_S_ and Gα_S_[Q227L].

STD-NMR experiments, combined with MD simulations, provided a window into the structure and dynamics of nucleotide interactions with Gα_S_ and Gα_S_ variants. STD-NMR data of Gα_S_ with GDP and GppNHp provided an interaction profile with which we could compare STD-NMR results of the studied variants. Compared with Gα_S_, for nearly all variants we observed a decrease in the STD-AF values for all observed protons with GDP-bound proteins (Figure 5 and 6). The exceptions were the variants Gα_S_[R265H] and Gα_S_[R258A], which both showed increased STD-AF values in complex with GDP. Both of these variants exhibited similar functional properties (Table S1) and similar thermal melting profiles (Figure 2), suggesting a potential correlation between changes in the nucleotide binding pocket with changes in global structure and function. MD simulations of selected disease-causing variants Gα_S_[Q227L], Gα_S_[R228C] and Gα_S_[R258A] showed that in all cases, although the residues that were mutated were not directly coordinating with the bound nucleotide, mutations disrupted the structure or dynamics of residues within the nucleotide binding pocket (Figure 7 and Figure S9). Although the resolution of the MD simulations was not sufficient to quantify significant variabilities of the average distance to the closest protein proton for each nucleotide proton, a more qualitative analysis could identify the closest residue in contact with each proton along the different MD trajectories, which could be evaluated with the context of the experimental data. The MD simulations were particularly insightful for revealing several unique features observed for Gα_S_[Q227L] (Figure 7 and Figure S9). For Gα_S_[Q227L], proton ‘c’, located on the pentose sugar ribose, exhibited significant differences in its local environment as compared with Gα_S_, Gα_S_[R228C], and Gα_S_[R258A]. Within the local environment of proton ‘c’, for Gα_S_[Q227L] we observed a significant increase in the percent contact with residue Glu50 (96.3%) and loss of contact with residues Asp173 and Arg201. Thus while Gα_S_[Q227L] shared some similarities in hydrogen bonding with Gα_S_, which may partially explain their similar thermal melting profiles, differences in the local nucleotide binding pocket dynamics may reflect different functions between Gα_S_[Q227L] and Gα_S_.

Upon comparing biophysical experimental and computational data across a range of Gα_S_ variants, we find no singular physical descriptor that consistently correlates with the impact of a mutation on Gα_S_ function. While mutations inducing similar functional alterations within proximal structural regions do share certain physical characteristics, such as similar thermal melting behavior, those with analogous functional changes do not consistently align with observations regarding global structural properties. In the more unique case of Gα_S_[Q227L], while the mutation appears to alter the local environment involving residues coordinating bound nucleotides, significant changes in the protein’s global conformation are not reflected in observations of the protein’s response to bound nucleotide. The comparison across all variants suggests that a uniform approach to designing molecules that may stabilize Gα_S_ variants or potentially partially restore function must account for their individual differences.

## Experimental procedures

### Generation of G*α*_S_ and variant G*α*_S_ plasmids

An initial Gα_S_ construct was gifted from the laboratory of Prof. Kevan Shokat (UCSF) (18), contained in a pET15b vector and derived from the short human isoform of Gα_S_ (PubMed accession number NP_536351) with the first 7 residues deleted. We modified this plasmid by replacing a DRICE cleavage site with a TEV cleavage site. This plasmid was used to express Gα_S_ for our experiments. All Gα_S_ variants were obtained from this plasmid by site-directed mutagenesis using the QuikChange II kit (Agilent) and primers listed in Table S2. The plasmid encoding for Mini-Gα_S_ was obtained from Genscript and designed to be identical to the sequence of the Mini-Gα_S_ ‘393’ sequence (25).

### Expression and purification of G*α*_S_ and G*α*_S_ variants

Each Gα_S_ plasmid was transformed into BL21-CodonPlus (DE3)-RIL cells (Agilent) and the bacteria were grown at 30 °C in M9 medium (50 mM Na_2_HPO_4_, 25 mM KH_2_PO_4_, 8.5 mM NaCl, 1 mM MgSO_4_, 50 µM FeCl_3_, 100 µM CaCl_2_, 44 mM Glucose, 56 mM NH_4_Cl and 0.01% w/v thiamine) with 34 µg/mL chloramphenicol and 100 µg/mL carbenicillin. Cells were grown to an optical density (OD_600nm_) between 0.6-0.8 and protein expression was then induced with 50 µM IPTG at 25 °C for 18-20 hours. Cells were sedimented by centrifugation at 4000 x g at 4 °C for 20 min then were lysed using a cell disruptor at 24,000 psi and 4°C in buffer containing 50 mL of 25 mM Tris pH 8.0, 150 mM NaCl, 1 mM MgCl_2_, 50 µM GDP with EDTA free in-house protease inhibitor cocktail (500 µM AEBSF, 1 µM E-64, 1µM leupeptin, 150 nM aprotinin) at a ratio of 50 mL lysis buffer for every 1 liter of cell culture. The lysate was clarified by centrifugation at 25,000 rpm at 10°C. The lysate was incubated overnight (16-20 hours) with 3 mL of NiNTA resin (Cytiva) for every 1L of cell culture at 4 °C. Then the resin was washed two times with 50 mL wash buffer containing 25 mM Tris pH 8.0, 500 mM NaCl, 1 mM MgCl_2_, 5 mM imidazole and 50 µM GDP at 4 °C. The first wash included 2 mg/mL iodoacetamide. The protein was eluted at 4 °C in a gravity flow column using a buffer containing 25 mM Tris pH 8.0, 250 mM NaCl, 1 mM MgCl_2_, 250 mM imidazole, 10% vol/vol glycerol, 50 µM GDP. The protein was then exchanged into a buffer containing 5 mM Tris pH 8.0, 100 mM NaCl, 10% vol/vol glycerol, 1 mM MgCl_2_ and 50 µM GDP via an Akta Start FPLC equipped with a HiPrep 26/10 desalting column (Cytiva). The protein was then incubated overnight at 4 °C with thermostabilized TEV protease (41) at a 1:50 molar ratio (Gα_S_:TEV). After overnight incubation (16-20 hours), cleaved Gα_S_ was purified via reverse immobilized mobile affinity chromatography (IMAC) by incubating the sample with nickel affinity resin at 4°C for 30 minutes and collecting the flow through using a gravity flow column. A final purification step was performed by size exclusion chromatography via an Akta Pure FPLC equipped with a Superdex 200 increase 10/300 GL column (Cytiva) equilibrated in buffer containing 20 mM HEPES pH 8.0, 150 mM NaCl, 5 mM MgCl_2_, 1 mM EDTA, 10% v/v glycerol, 10 µM GDP. Peak fractions containing purified Gα_S_ or Gα_S_ variants were isolated and pooled for subsequent experiments. 10% SDS gels were prepared to check the purity and confirm the identity of the protein. The gel was made with running buffer consisting of 1 M Tris pH 8.4, 10% acrylamide (BIORAD), 0.1% SDS, 0.12% APS, 8 mM TEMED (BIORAD) and stacking buffer consisting of 0.75 M Tris pH 8.4, 5% acrylamide (BIORAD), 0.1% SDS, 0.1% APS, 7 mM TEMED (BIORAD) and run from 10X stock of cathode buffer (1 M Tris pH 8.4, 1M tricine, 1% SDS) and 10X anode buffer (2 M Tris pH 8.9) diluted to 1X. Apo Gα_S_ proteins and variants (without nucleotide) were purified in the same manner as above but without GDP in any of the buffers.

### Expression and purification of mini-G*α*_S_

We expressed and purified mini-Gα_S_ using a strategy adapted from earlier studies (25,26). BL21-CodonPlus (DE3)-RIL were transformed with the expression vectors and grown at 30 °C in Terrific broth (TB) medium with 0.2% glucose, 34 µg/mL chloramphenicol, 100 µg/mL carbenicillin and 5 mM MgSO_4_. Protein expression was induced with 50 µM IPTG at 25 °C and left to grow overnight (16-20 hours). Cells were lysed in 40 mM HEPES pH 7.5, 100 mM NaCl, 10 mM imidazole, 10% vol/vol glycerol, 5 mM MgCl_2_, 50 µM GDP with EDTA free in-house protease inhibitor cocktail (500 µM AEBSF, 1 µM E-64, 1 µM leupeptin, 150 nM aprotinin) and using a cell disruptor operating at 24 kPsi and 4°C. The lysate was clarified by centrifugation at 25,000 rpm and 10°C for 30 min and loaded on a His-Trap HP NiNTA 5 mL column (Cytiva) via an Akta Start FPLC. Resin was washed with lysate buffer containing a final concentration of 40 mM imidazole and 500 mM NaCl and protein was eluted from the resin with lysate buffer containing 400 mM imidazole. The sample was exchanged into buffer via an Akta Start FPLC equipped with a HiPrep 26/10 desalting column (Cytiva) that was equilibrated in 20 mM HEPES pH 7.5, 100 mM NaCl, 10% vol/vol glycerol, 1 mM MgCl_2_, 10 μM GDP. Fractions containing mini-Gα_S_ were collected and incubated for 16-20 hours at 4 °C with thermostabilized TEV protease at 1:100 molar ratio (mini-Gα_S_:TEV. Cleaved mini-Gα_S_ was purified via reverse IMAC by incubating the sample with nickel resin for 30 minutes and collecting the flow through using a gravity flow column. Mini-Gα_S_ was further purified via an Akta Start FPLC equipped with a HiLoad Superdex 75 pg column (Cytiva) equilibrated in storage buffer (10 mM HEPES pH 7.5, 100 mM NaCl, 10% vol/vol glycerol, 1 mM MgCl_2_, 1 μM GDP, 0.1 mM TCEP). Purified samples were rapidly frozen in liquid nitrogen and stored at −80 °C until needed for experiments.

### Tryptophan fluorescence assays

Gα_S_ and Gα_S_ variants were exchanged into buffer containing 20 mM HEPES pH 8.0, 150 mM NaCl, and 5 µM GDP using a PD MiniTrap G25 column (Cytiva), and the sample volumes were adjusted to achieve a final concentration of 10 µM protein. Tryptophan fluorescence spectra were recorded with a Cary Eclipse spectrometer operating at 25 °C. The excitation wavelength was 280 nm and emission was observed at 340 nm with 5 nm excitation and emission bandwidths. GTPγS was added to 200 µL protein for a final concentration of 50 µM in a quartz cuvette. The sample was mixed vigorously by repeated pipetting and data collection immediately started. Data points were recorded every 12 seconds for 200 minutes total acquisition time. Data were analyzed in GraphPad prism using the one-phase association equation shown below:

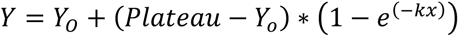

where *Y_O_* is the fluorescence intensity when *x* (time) equals zero, ‘Plateau’ is the fluorescence intensity when the system has been fully equilibrated at infinite time expressed in the same units as *Y*, and *k* is the rate constant, expressed as the reciprocal of the x-axis time units.

### Protein thermal unfolding monitored by circular dichroism

Single-wavelength variable temperature circular dichroism data were recorded on a Chirascan qCD spectrometer. Gα_S_ or Gα_S_ variants were concentrated to 10 µM in buffer containing 10 mM HEPES pH 7.5, 100 mM NaCl, 1 mM MgCl_2_, and 50 µM GDP or 50 µM GTPγS. First, full-wavelength CD spectra were recorded at 25 °C to confirm samples were properly folded. Then, protein melting was monitored by recording the ellipticity at 220 nm from 25 °C to 89 °C with a linear heating rate of 1 °C/min. Each melting temperature (T_m_) was determined by fitting the data in GraphPad prism to a Boltzmann sigmoidal equation shown below:

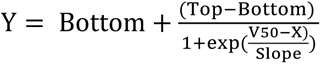

Where *Y* is the fraction of folded protein as a function of temperature, V_50_ is the temperature at which half of the protein is unfolded, X represents the temperature, and ‘Slope’ is the steepness of the curve. ‘Top’ and ‘Bottom’ are normalization factors set to 1.0 and 0, respectively. Error bars were determined by calculating the standard deviation from three independent measurements.

### NMR experiments

Samples of Gα_S_ or Gα_S_ variants were exchanged into NMR buffer using a PD MiniTrap G25 column (Cytiva) equilibrated with 25 mM HEPES pH 7.0, 75 mM NaCl, 5 mM MgCl_2_, 100 µM DSS, 10 % v/v ^2^H_2_O, and either 2 mM GDP or 2 mM GppNHp. Exploratory experiments employing a range of protein-to-ligand concentrations were tested to determine optimal STD-NMR signal-to-noise. Based on these data we prepared samples containing 40 µM protein and 2 mM ligands (Figure S6). NMR experiments were recorded at 25°C and 800 MHz ^1^H Larmor frequency using a Bruker Avance III spectrometer equipped with a 5 mm TXI cryoprobe. 1D-^1^H saturation transfer difference (STD) experiments were recorded using water suppression by gradient-based excitation sculpting (Brucker pulse program stddiffesgp), with 64 scans per experiment and a 3 s delay between scans. On-resonance spectra were acquired by irradiating at 0.79 ppm to target Gα_S_ methyl protons, and off-resonance were acquired by irradiating at −40.0 ppm. Saturation times were tested from 0.5 to 3 s, and an optimal saturation time of 2 s was determined (Figure S5). Data were acquired and analyzed in Topspin 4.1.1. Prior to Fourier transformation, the data set was multiplied by an exponential window function applying 3 Hz line broadening and zero-filled to 65536 points; baseline correction was applied to all frequency domain data sets. To reduce background signals, a reference STD-NMR spectrum was recorded with a sample containing the same concentration of protein but no nucleotide. This spectrum was processed identically and subtracted from all STD-NMR spectra.

Amplification factors, A_STD_, were determined from the integrals of ligand signals in the STD difference (I_0_ − I_sat_) and reference (I_0_) spectra according to the following equation from reference (31)

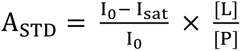

where A_STD_ is the amplification factor of each STD signal, I_O_ is the integral of the STD signal in the reference spectrum, I_sat_ is the integral of the STD signal in the difference spectrum and [L] and [P] are the concentrations of ligand and protein, respectively.

Signal-to-noise ratios in the STD difference spectra, *SNR_diff_*, were determined for each ligand signal using the “sinocal” function on topspin 4.1.1. Error associated with the amplification factors, σ, was evaluated using the following equation:

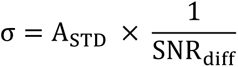

where A_STD_ is the amplification factor of each STD signal and SNR_diff_ is the signal to noise ratio of the STD signal. For presentation of the normalized A_STD_ signals, each nucleotide signal “a” to “d” and associated errors were normalized by the *A_STD_* of the “a” signal.

### Molecular Dynamics Simulations

The crystal structure of the Gα_S_ protein in complex with GTPγS was retrieved from the Protein Data Bank (PDB ID 1AZT) (42), and prepared for MD simulations as follows. Chain B from the dimer in the asymmetric unit was retained, and the missing segment between α1 and αA (residues 70-86) were modeled with Prime (Schrödinger Release 2011-2: Prime; Schrödinger, LLC: New York, 2011). Crystallographic water molecules beyond 5 Å of GTPγS as well as all co-crystallized phosphate ions were removed, and the protonation states of ionizable residues were determined with PROPKA at pH 7.0. The GTPγS substrate was modified to either GTP or GDP, depending on the intended system, with Maestro v 9.4 (Schrödinger, LLC; NY). Amino acid replacements to generate the different Gα_S_ variants were produced with the Schrödinger’s PyMOL mutagenesis tool (The PyMOL Molecular Graphics System, Version 2.0. Schrödinger, LLC.)

MD simulations were executed with the Q software package (43), using the OPLS-AA/M force field for proteins and the TIP3 water model (44). The parameters to describe GTP and GDP were taken from same force field. All models were partially solvated in a water sphere of diameter 50 Å with its center located at the nucleotide’s center of geometry. Protein atoms outside the sphere were tightly constrained to their initial coordinates with a force constant of 200 kcal mol^-1^ Å^-1^ and excluded from nonbonded interactions, while water molecules at the sphere boundary were subjected to radial and polarization restrains following the SCAAS model (45). All atoms inside the simulation sphere were allowed to move freely, where the local reaction field multipole expansion was used for long-range electrostatic interactions beyond a cut-off of 10 Å (46). Each system was subject to 100 independent replica MD simulations, each of them starting with an equilibration phase consisting of 51 ps of gradual heating to 298 K and concurrent release of (initial 10.0 kcal mol^-1^ Å^-2^) harmonic restraints on solute heavy atoms. Unrestrained MD simulations followed for 5 ns at a temperature of 298 K and employed a 1 fs time step, giving a total of 0.5 μs of unrestrained simulation per system, accumulating 2 μs MD simulations in this study.

MD analyses included pairwise contacts, backbone RMSF and H-bond analysis based on cutoffs for the Donor-H···Acceptor distance (< 2.4 Å) and angle (> 120 deg), was performed with the MDtraj open library (47). All structural images were generated with PyMOL.

## Data availability

All data supporting the findings of this study are available within the paper and supporting information. Unprocessed NMR data will be made available upon request. Correspondence

## Supporting information

This article contains supporting information.

## Acknowledgements

The authors would like to thank Dr. James Collins and Mr. James Rocca from the UF AMRIS facility for providing training and assistance with NMR experiments.

## Authors Contributions

G.F., K.A., and M.T.E. conceptualization; K.A., G.F., E.P., L.K., H.G.d.T. and M.T.E. investigation; K.A., G.F., L.K., H.G.d.T. and M.T.E. formal analysis; K.A., G.F., L.K., H.G.d.T. and M.T.E. writing original draft; L.K., K.A., H.G.d.T and M.T.E. writing-reviewing and editing; H.G.d.T and M.T.E. project administration.

## Funding and additional information

We acknowledge the National Institutes of Health grant number R35GM138291 (G.F., M.T.E., K.A., and E.P.) and a University of Florida Graduate student fellowship (K.A.) for support on this work. A portion of this work was supported by the McKnight Brain Institute at the National High Magnetic Field Laboratory’s AMRIS Facility, which is funded by National Science Foundation Cooperative Agreement No. DMR-1644779 and the State of Florida. The computations were enabled by resources provided by the National Academic Infrastructure for Supercomputing in Sweden (NAISS) and the Swedish National Infrastructure for Computing (SNIC) partially funded by the Swedish Research Council through grant agreements no. 2022-06725 and no. 2018-05973.

## Abbreviations

CD: Circular dichroism
NMR spectroscopy: nuclear magnetic resonance spectroscopy
STD-NMR: saturation-transfer difference-NMR
GPCR: G protein-coupled receptor
MD: molecular dynamics

## Supporting information

### Supplementary results

**Figure S1.**
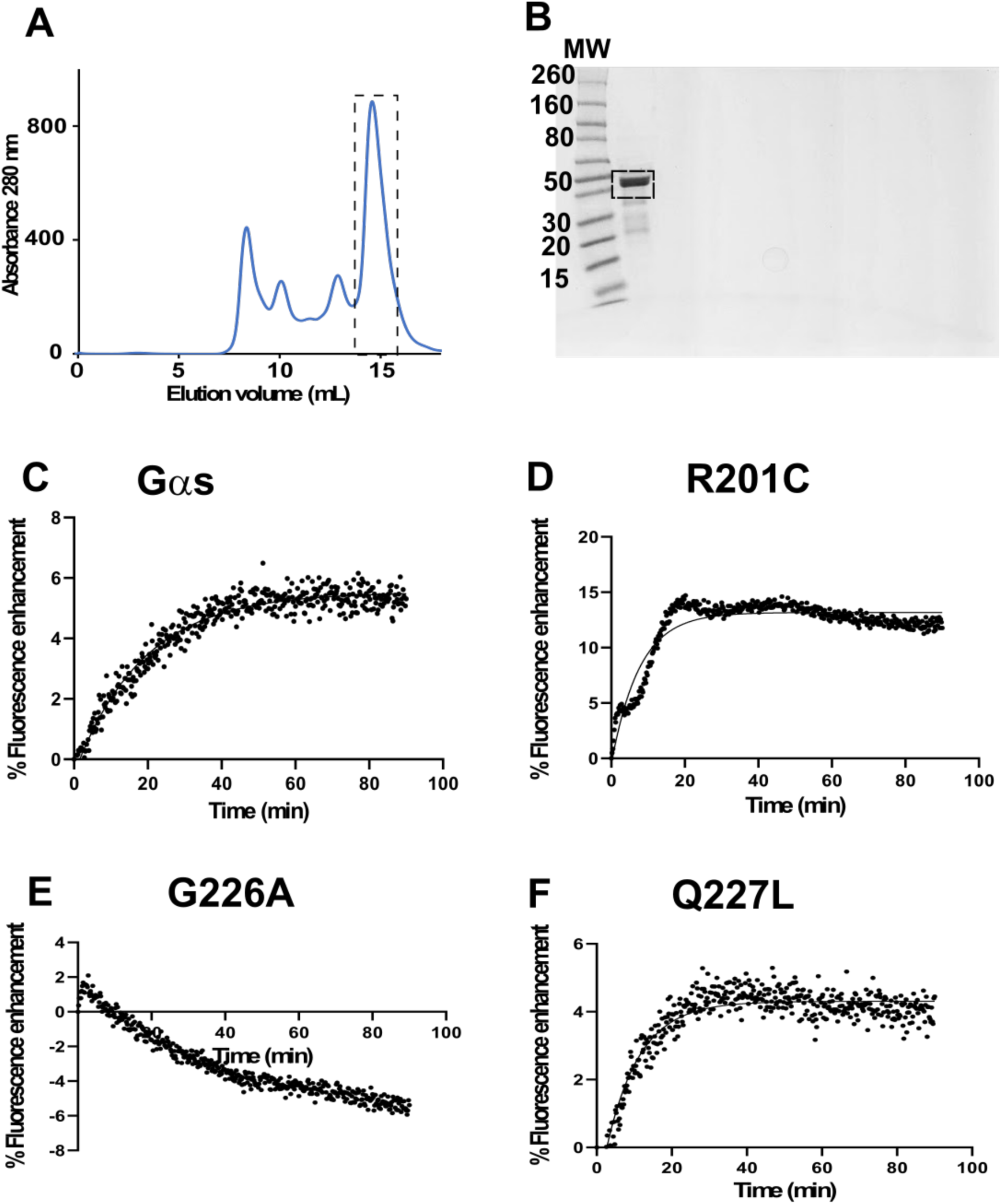
Purification of Gα_S_ protein and functionality. *A*, size exclusion chromatogram of Gα_S_ protein purification shown with the boxed peak corresponding to the Gα_S_ protein. *B,* SDS page gel of purified Gα_S_ protein with the molecular weight standard indicating the size of the 44kDa protein. *C,* tryptophan fluorescence assay of Gα_S_ protein showing an increase in tryptophan fluorescence upon GTP binding and G protein activation *D,* tryptophan fluorescence assay of the hyperactivating R201C variant showing a faster increase in tryptophan fluorescence upon GTP binding. *E,* tryptophan fluorescence assay of G226A variant showing no increase in tryptophan fluorescence upon GTP binding. *F,* tryptophan fluorescence assay of the hyperactivating Q227L variant showing a faster increase in tryptophan fluorescence upon GTP binding.

**Table S1.**
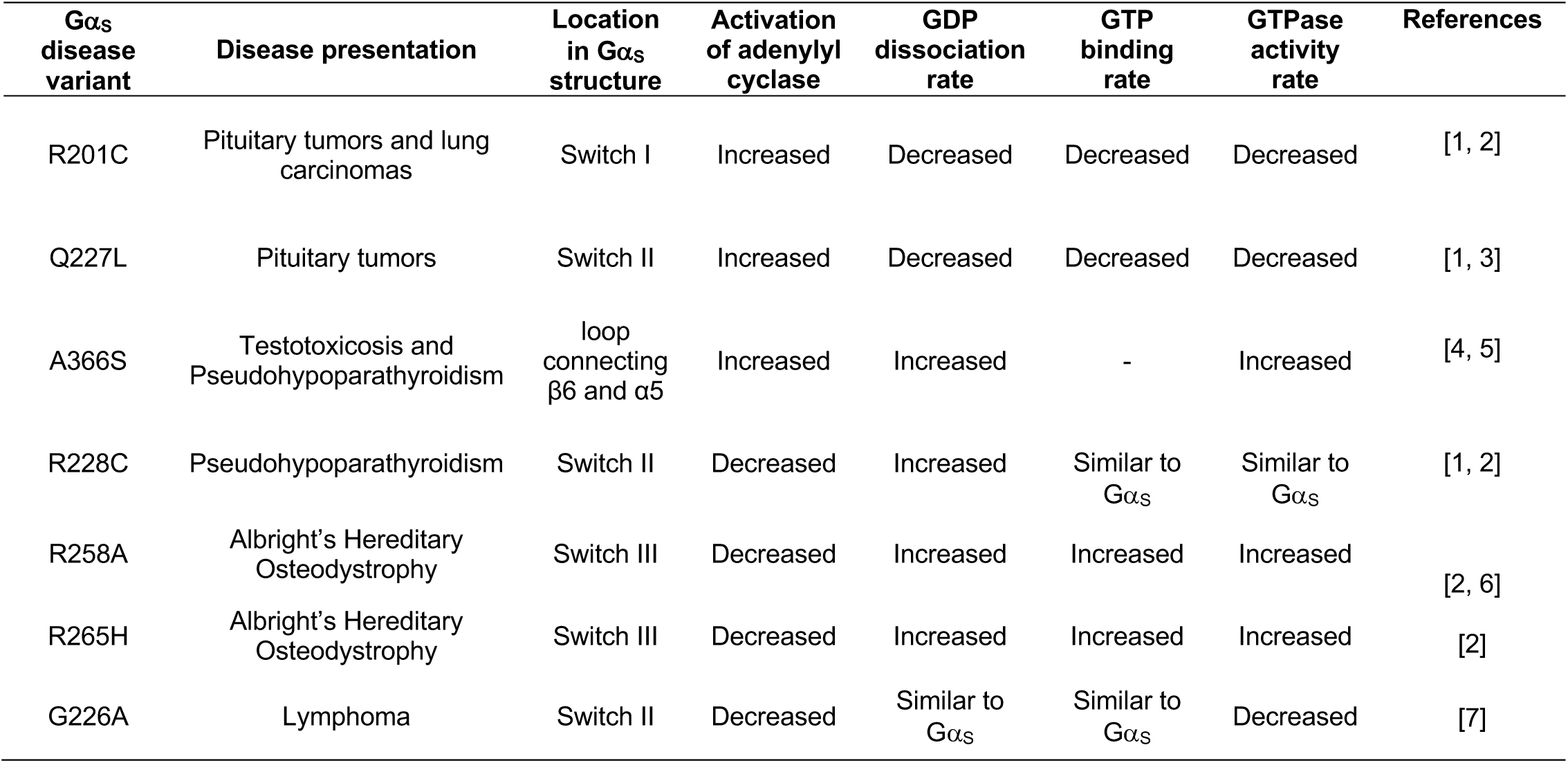
Summary of biochemical properties of Gα_S_ disease causing variants. Each variant is characterized in terms of which disease state it is represented in, the location in the Gα_S_ structure, the effect of the mutation on the activation of adenylyl cyclase, the rate of GDP dissociation, the rate of GTP binding and the rate of GTPase activity. The dash indicates that this particular value was not determined within the cited study.

**Table 2.**
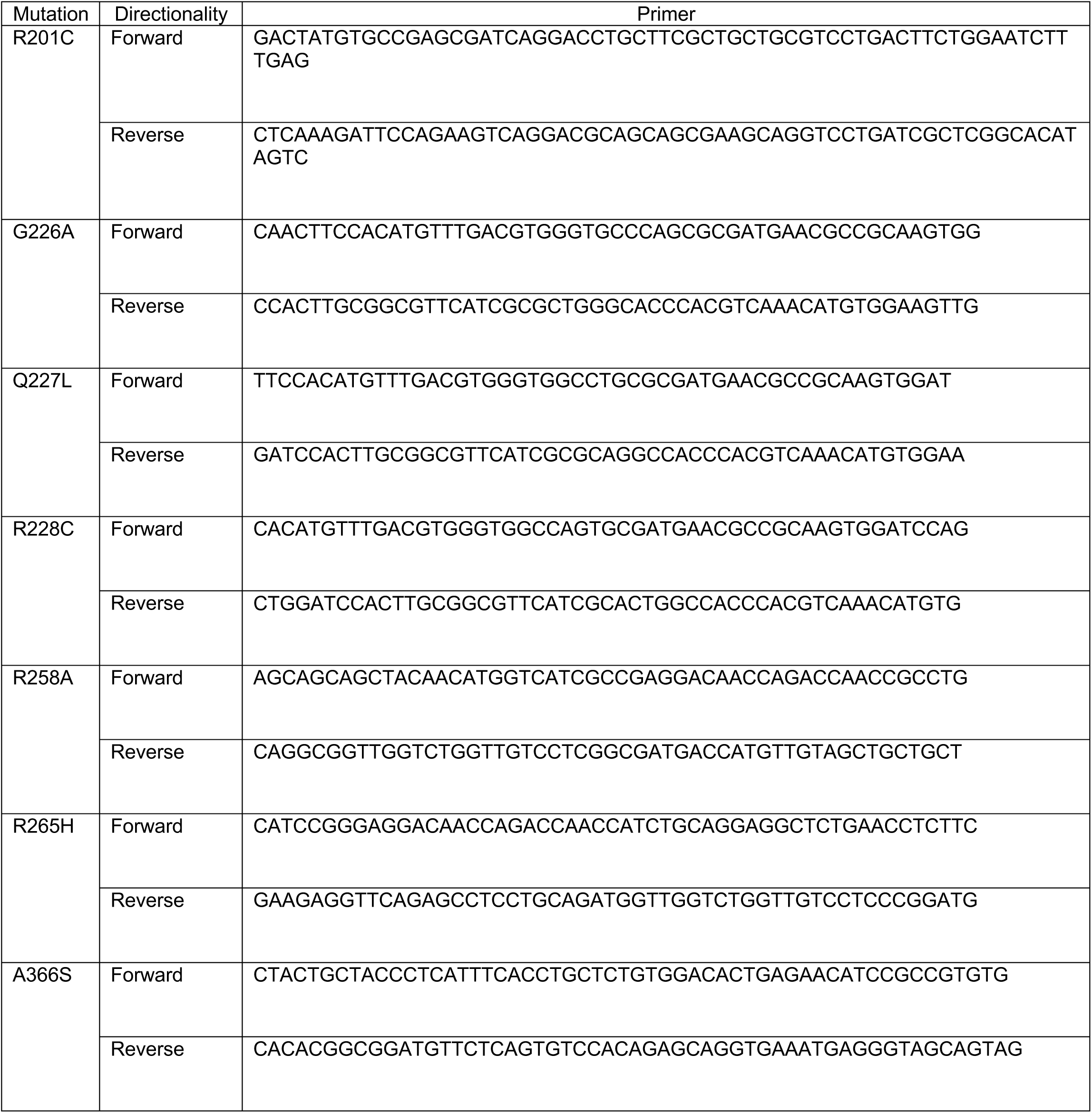
Primers designed for Gα_S_ variants. List of forward and reverse primers designed to obtain Gα_S_ disease-associated variants.

**Figure S2.**
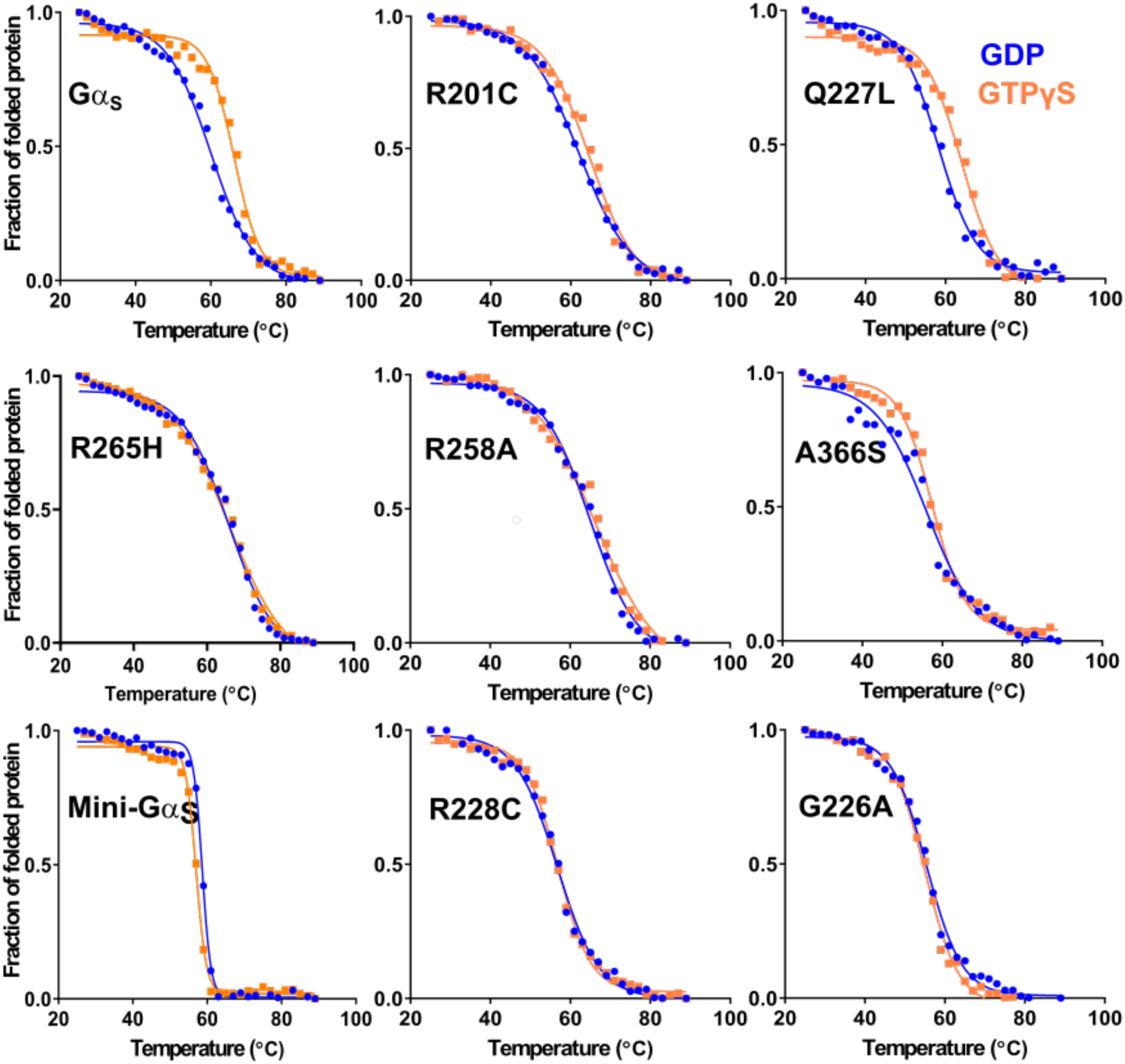
Thermal melting profile of Gα_S_ and diseased variants in GDP and GTPγS determined by circular dichroism. The thermal unfolding of Gα_S_ and disease-associated variants bound to GDP and GTPγS monitored by variable temperature single wavelength CD. Same color scheme as Figure 1*A*. The thermal melting temperatures were obtained by fitting the data to a Boltzmann sigmoidal function in GraphPad prism.

**Figure S3.**
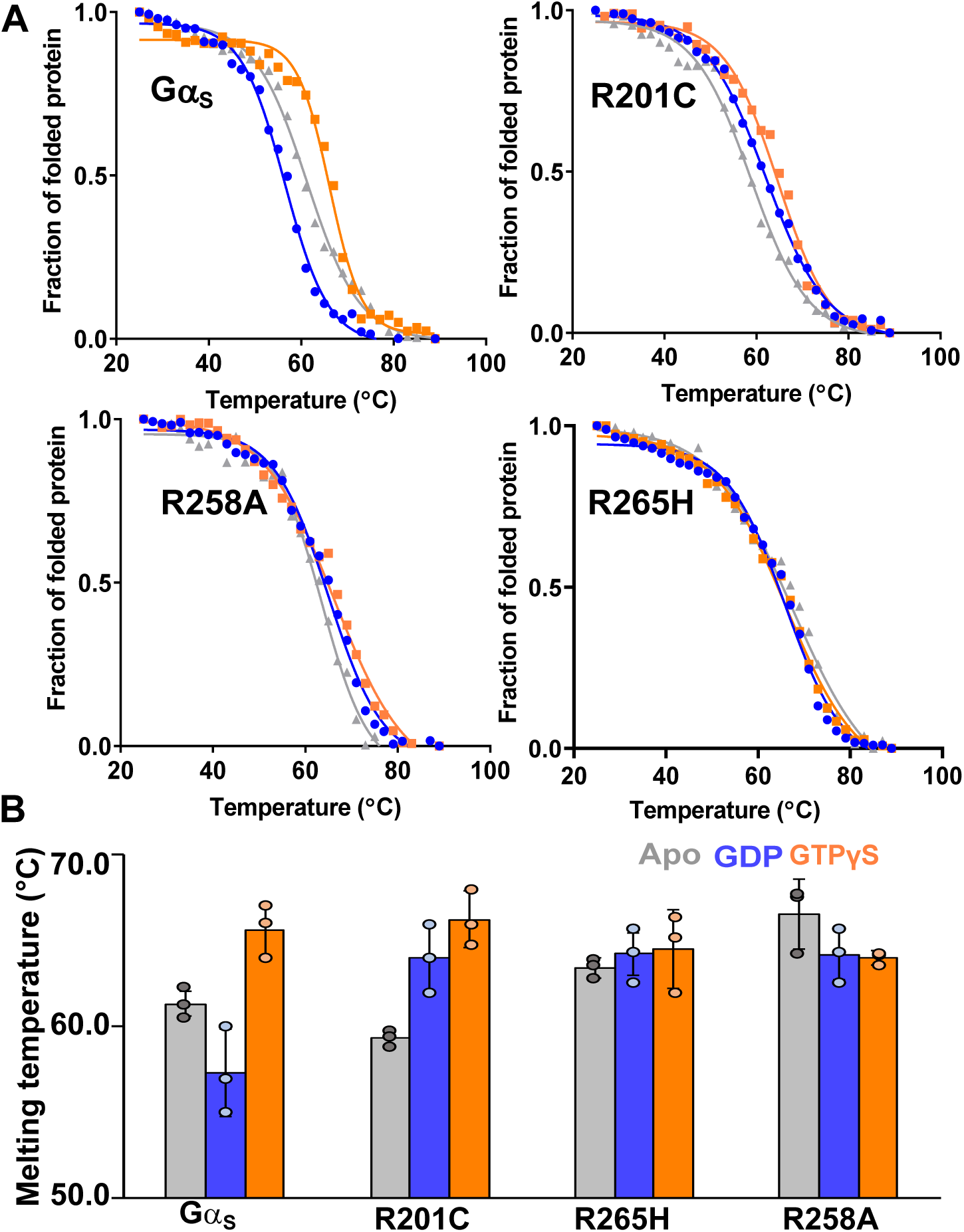
Thermal melting profiles of Gα_S_ and Gα_S_ variants in the presence of GDP, GTPγS, or with no nucleotide added (apo). *A,* The thermal unfolding of Gα_S_ and variants Gα_S_[R201C], Gα_S_[R258A] and Gα_S_[R265H] when bound to GDP, GTPγS or with no nucleotide added (apo), as monitored by variable temperature single wavelength CD. Same color scheme used as in Figure 1. *B,* histograms of the melting temperature (T_m_) values determined by fitting the data shown in panel A. Error bars represent the standard deviation of triplicate measurements.

**Figure S4.**
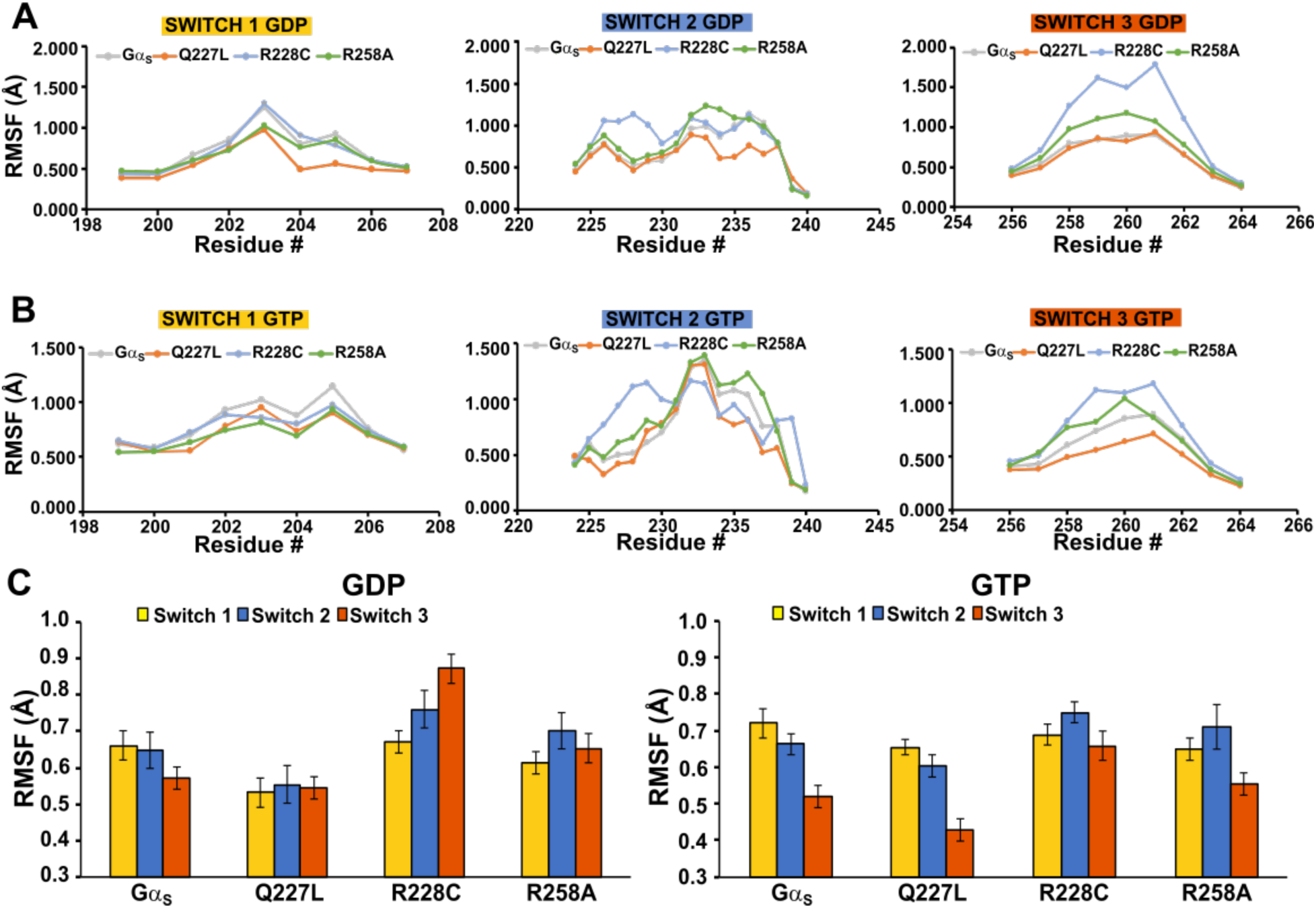
Backbone root mean square fluctuations (RMSF) values within the switch regions of Gα_S_ and Gα_S_ variants in complexes with GDP or GTP. *A* and *B,* line plots of backbone RMSF values of residues in Switch I, II and III regions for Gα_S_ (gray), Gα_S_[Q227L] (orange), Gα_S_[R228C] (light blue) and Gα_S_[R258A] (green). *C,* histograms of the average RMSF values for Gα_S_ and the Gα_S_ variants bound to GDP (left panel) and GTP (right panel) in the regions of Switch I (yellow bars), Switch II (blue bars) and Switch III (red bars). Error bars in the histograms represent the standard error of the mean.

**Figure S5.**
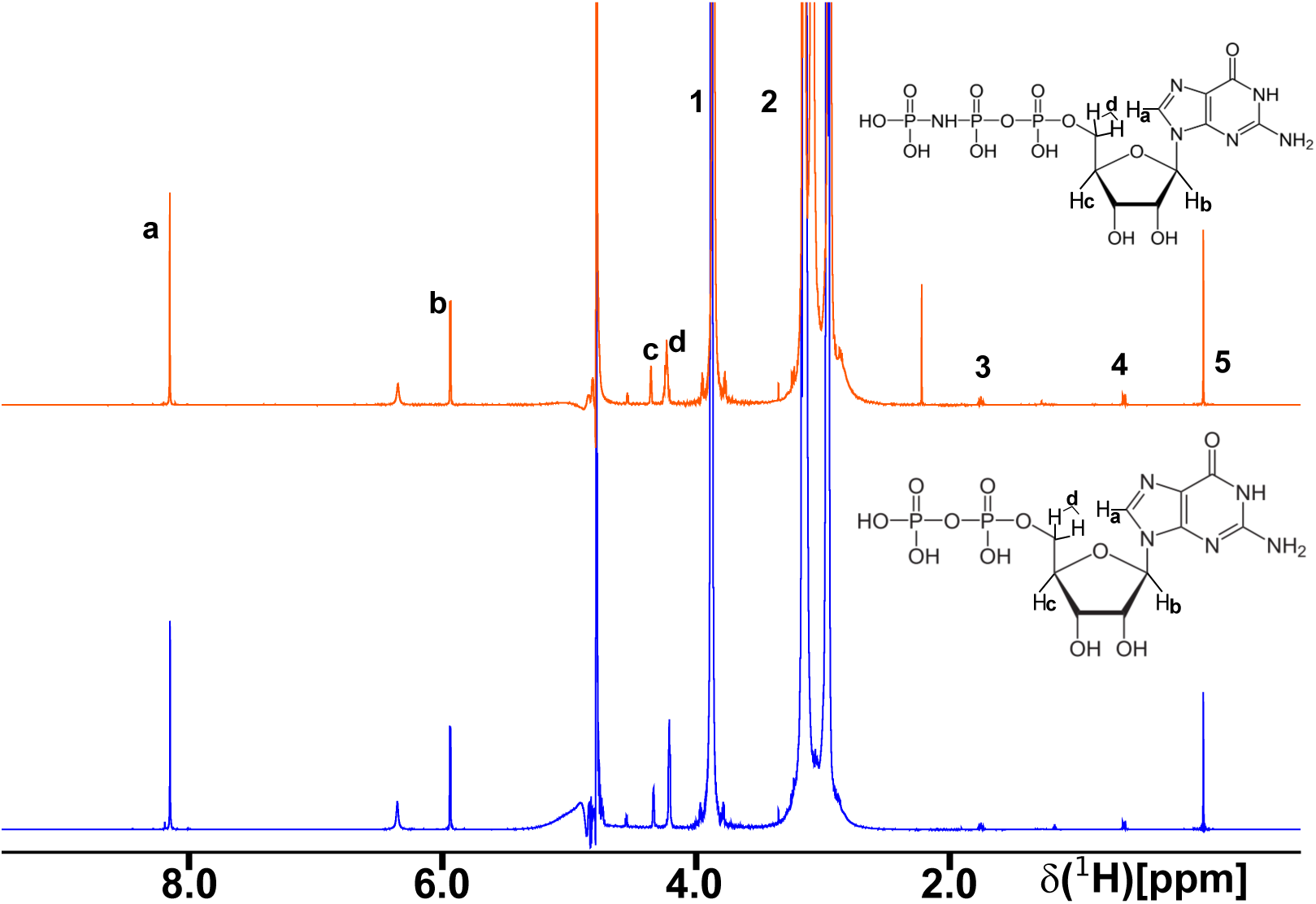
^1^H signal assignment in reference spectra of GDP and GppNHp. 1-dimensional ^1^H-NMR reference spectrum of GDP (blue) and GppNHp (orange) nucleotide. ^1^H signals labeled ‘a’ through ‘d’ were utilized in STD-NMR experiments and are shown on the chemical structures of GDP and GppNHp. Assignments were transferred from BMRB entry bmse000270. ^1^H signals labeled 1 and 2 are from HEPES buffer, and ^1^H signals labeled 3-5 are from the sodium trimethylsilylpropanesulfonate (DSS) NMR standard.

**Figure S6.**
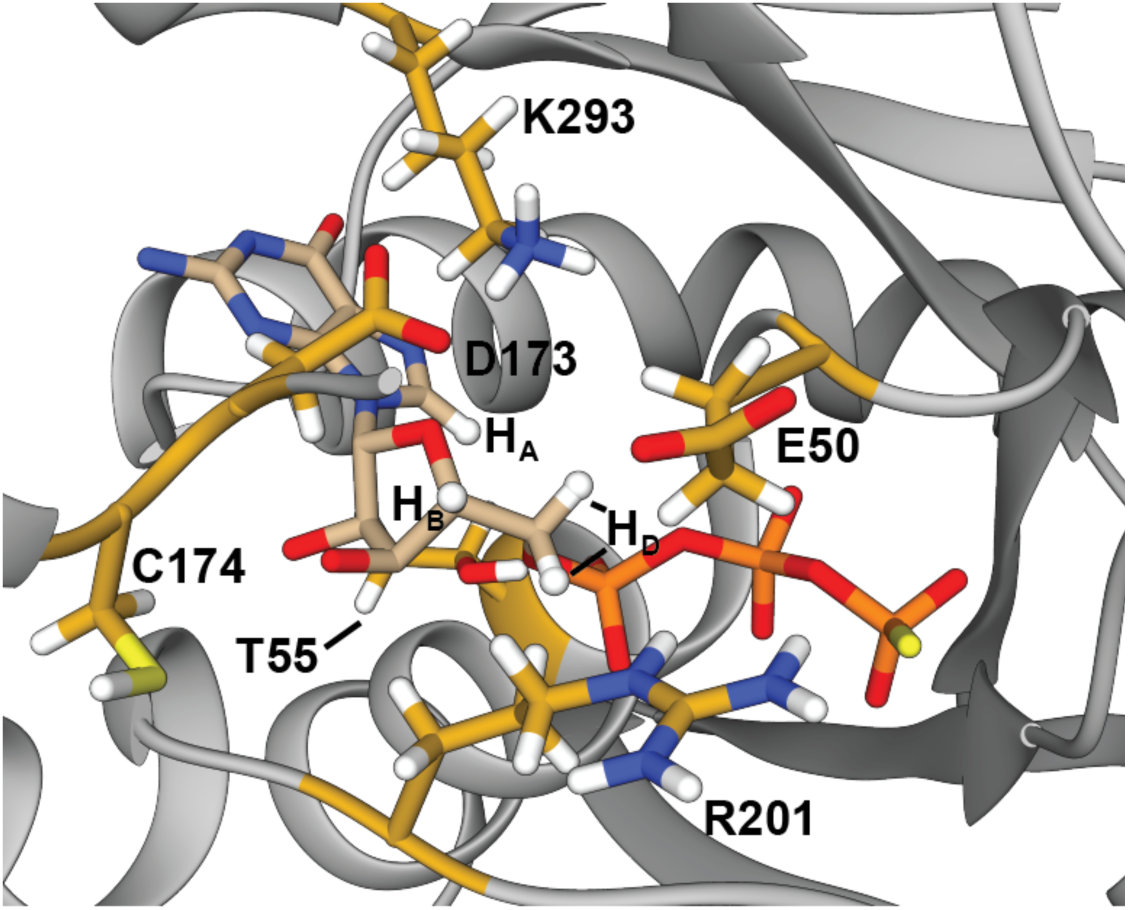
Residues closest to bound GTP in the Gα_S_ nucleotide binding pocket. An expanded view is shown of the Gα_S_ nucleotide binding pocket with GTP bound. (shown in tan stick representation, PDB:1AZT) with protons seen in STD-NMR interactions annotated ‘H_A_’ through ‘H_D_’. Residues within 6Å of these protons are annotated.

**Figure S7.**
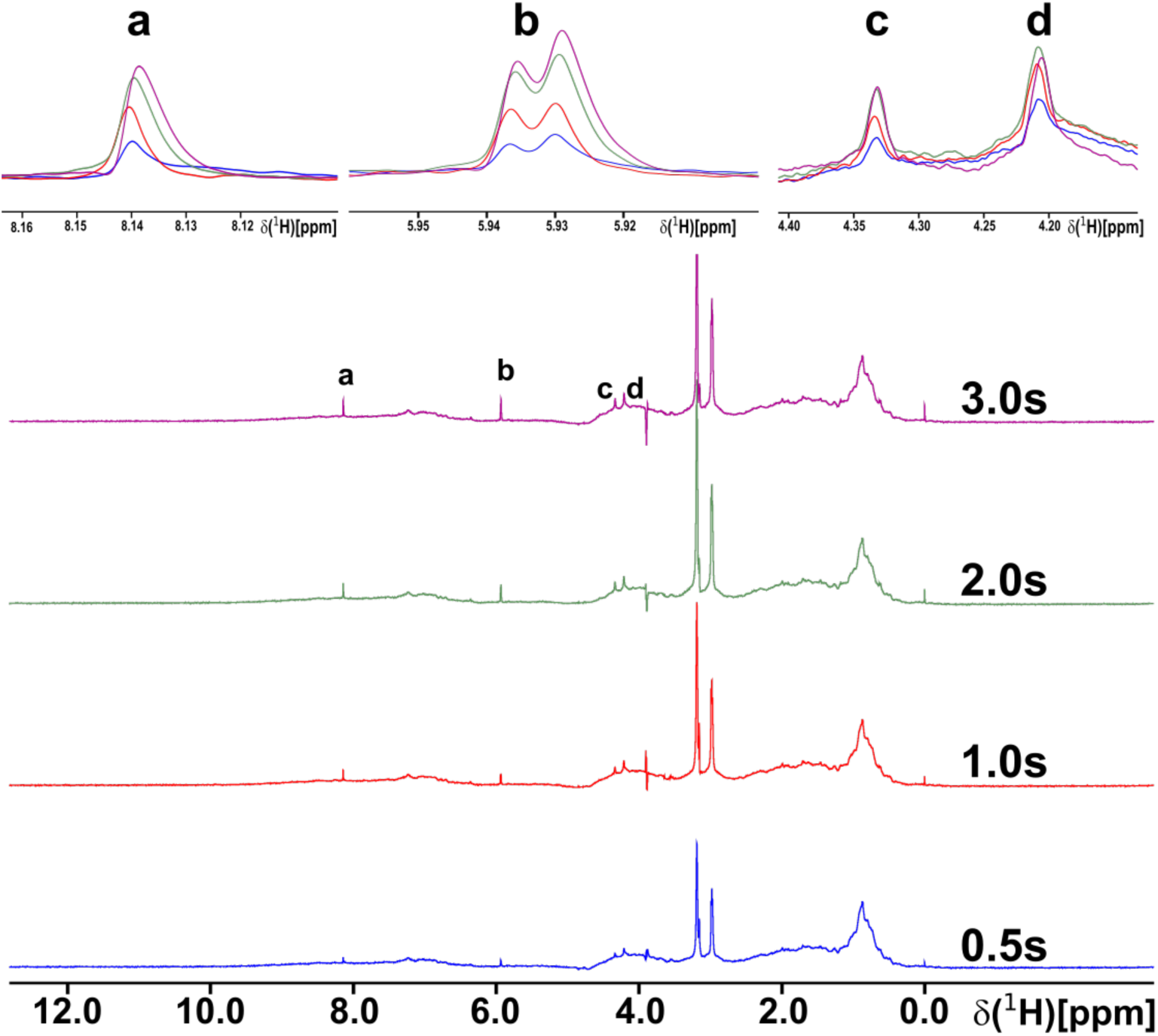
Optimization of STD-NMR saturation transfer time. ^1^H STD-NMR spectra are shown with 40 µM Gα_S_ and 2 mM GDP recorded with four different saturation transfer times between 0.5 s and 3.0 s. ^1^H signals labeled ‘a’ through ‘d’ were used to calculate STD-NMR amplification factors. Expanded views of each signal are shown in the panels at the top.

**Figure S8.**
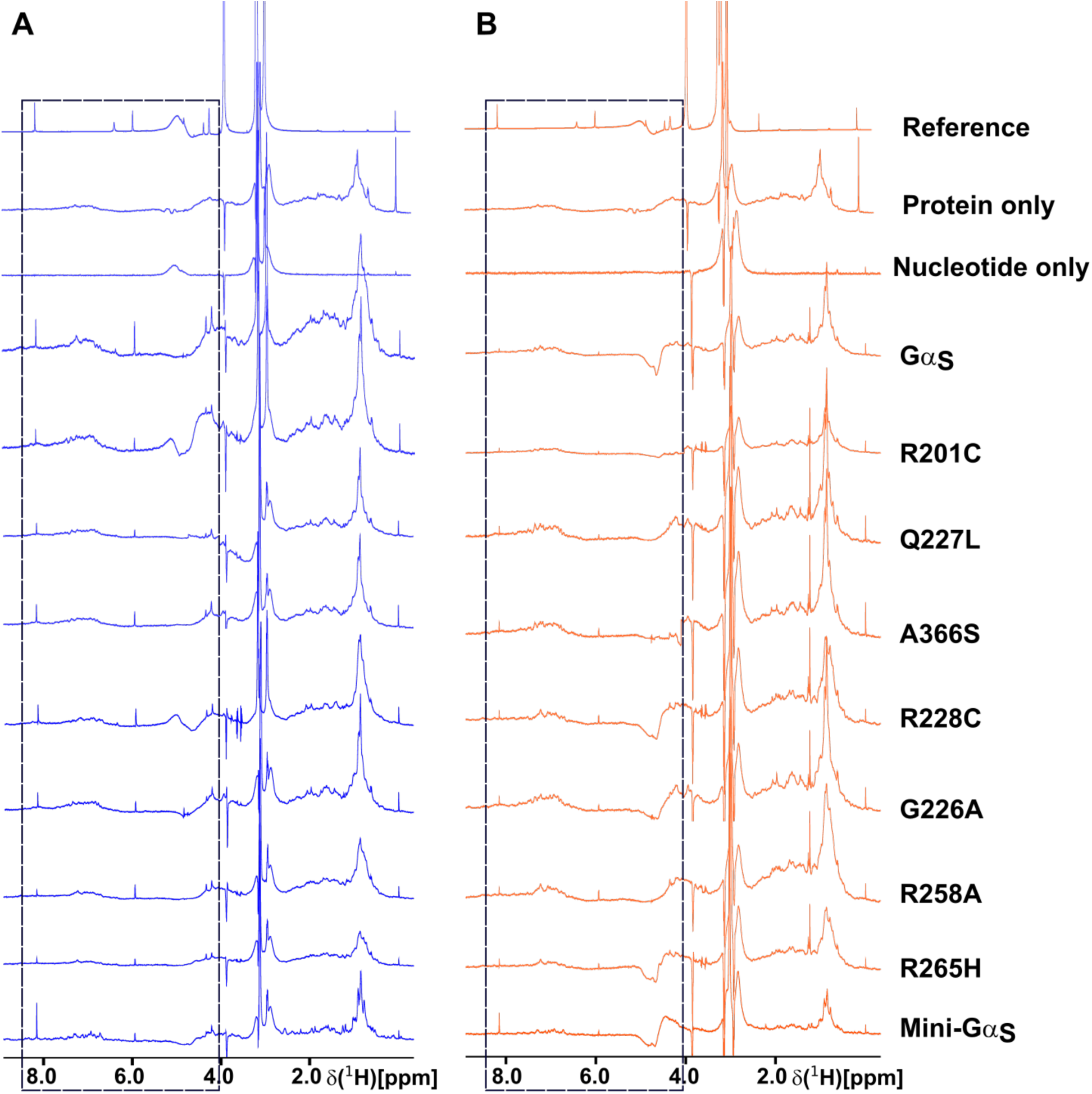
One-dimensional ^1^H STD-NMR spectra of Gα_S_ and Gα_S_ variants in complexes with GDP and GppNHp. STD-NMR spectra of Gα_S_ and variants in complex with GDP shown in panel *A* (in blue) and GppNHp shown in panel *B* (in orange). “Reference” is a 1D ^1^H NMR spectrum of GDP, “protein only” is a STD-NMR control experiment with a sample containing Gα_S_ and buffer but no nucleotide, and “nucleotide only” is a STD-NMR control experiment with a sample containing nucleotide and buffer but no protein. Boxed regions indicate areas of interest where an STD-NMR effect is seen.

**Figure S9.**
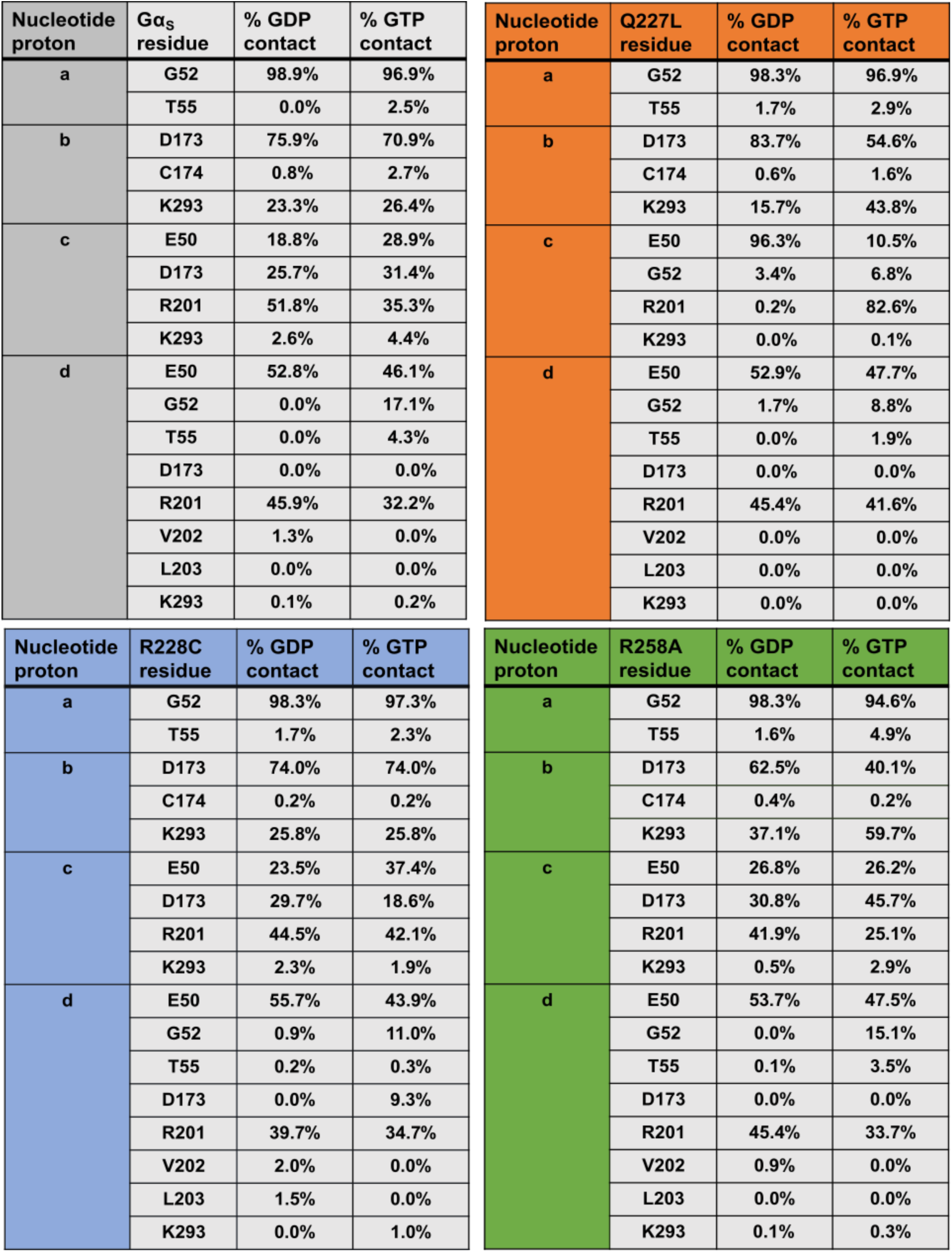
Percent of contact of residues of Gα_S_ and Gα_S_ variants with GDP and GTP protons in STD-NMR over MD simulations. The percent contact of residues of the Gα_S_ protein and Gα_S_[Q227L], Gα_S_[R228C] and Gα_S_[R258A] in contact with the protons “a” to d” (d is the average of chemically equivalent protons) in GDP and GTP over the course of MD simulations.

